# Active Merlin Binds RalB to Regulate Exocytosis

**DOI:** 10.1101/2025.06.13.659557

**Authors:** Robert F. Hennigan, Kayley McLaughlin, Nancy Ratner

## Abstract

Loss of *NF2* tumor suppressor activity causes NF2-related schwannomatosis. Proximity biotinylation identified proteins proximal to Merlin isoform 1 and isoform 2 at confluence, when Merlin is active, but not in sub-confluent, growing cells. These data confirmed Merlin involvement in cell-cell and cell-substrate junctions, identified new signal transduction pathways, and highlighted a role for Merlin in intracellular transport. Direct binding assays identified the small GTPases RalA and RalB as high affinity PIP_2_-dependent Merlin binding proteins that co-localized with RalA/B on the plasma membrane. Merlin loss resulted in aberrant activation of RalA and RalB at high cell density. Merlin competitively inhibited RalB binding to its exocyst effectors Sec5 and Exo84 and regulated the kinetics of exocytosis in a RalB dependent manner. Thus, RalB is a novel binding partner for active Merlin, and the RalA/B pathway is a possible therapeutic target to treat NF2-related schwannoma.

## Introduction

NF2-related schwannomatosis, formerly called Neurofibromatosis type 2, is an inherited disease characterized by benign peripheral nerve tumors known as schwannomas, slow growing tumors that cause significant morbidity and are resistant to chemotherapy^1^. *NF2*-mutant schwannomas also arise in the general population, in which they represent 8% of all intracranial tumors^2,3^. *NF2* loss is also common in meningiomas, mesotheliomas, and other types of sporadic tumors. Targeted deletion of the *Nf2* gene in mouse Schwann cells leads to schwannoma^4^*, c*onfirming that *NF2* is a bona-fide tumor suppressor gene, with loss of function causing tumorigenesis. The major phenotype of *Nf2*-null cells *in vitro* is impaired contact inhibition of growth^5^, a fundamental hallmark of cancer and a key feature of NF2 pathogenesis.

The *NF2* gene encodes Merlin, a 70-kDa member of the Ezrin-Radixin-Moesin (ERM) branch of the band 4.1 superfamily^6^. Merlin secondary structure consists of an N-terminal FERM domain followed by a central α-helical region that positions the C-terminal domain (CTD) for intramolecular interaction with the FERM domain^7^. Upon the release of the CTD from the FERM domain, Merlin transitions to a more open “active” conformation that allows Merlin dimerization and FERM domain interaction with critical binding proteins^8^. A Merlin mutant that stabilizes the FERM-CTD interaction adopts a closed conformation that has impaired tumor suppressor activity^8^. Conversely, a more open, FERM-accessible conformation mutant retains tumor suppressor activity^8^. Merlin is active when cells are contact inhibited at confluence and inactive in growing, sub-confluent cells^5^. Merlin has two major splice variants, isoform 1 and isoform 2, that differ in their extreme C-terminal amino acids^9^. The isoform 2 C-terminus has a lower affinity for the FERM domain^10^, a more open conformation^8^ and increased ability to dimerize^11^. However, both Merlin isoform 1 or isoform 2 are tumor suppressors^12^ and mutations in both isoforms are detected in tumors^9^.

Merlin is localized predominately to the inner face of the plasma membrane^13^ where it is associated with a variety of cell to cell and cell to substrate junctional complexes, including integrin-based focal adhesions and cadherin-based adherens junctions^14^. This localization is facilitated by three pairs of basic amnio acids in the FERM domain that are necessary for association with lipid rafts and for growth suppression^15^. Membrane localization positions Merlin for binding the lipid second messenger phosphotydylinositol-4,5-biphosphate (PIP_2_). PIP_2_ binding causes an allosteric change in the Merlin central α-helical domain that forces the CTD and FERM domains apart^16^, allowing Merlin to assume it’s open, FERM accessible conformation^17^. PIP_2_-mediated transition to an open conformation leads to Merlin dimerization and increased affinity for target proteins including Lats1, YAP1 and ASPP2^11,16^. These data support a model in which Merlin is activated at high cell density by dimerization in response to transient, localized increases in PIP_2_ levels, leading to contact inhibition of growth.

In mammalian cells, contact inhibition of growth is mediated by the Hippo pathway, a growth inhibitory kinase cascade that responds to mechanosensory cues transmitted via cell junctional signaling complexes^18^. Merlin regulates the HIPPO pathway by binding to and activating the core HIPPO kinase, Lats1/2. Lats1/2 then phosphorylates the transcriptional co-activator YAP1, leading to its ubiquitination and degradation, thereby preventing YAP1 nuclear localization, leading to growth arrest^19^. While impaired Hippo signaling due to Merlin loss is considered a primary mechanism contributing to NF2 schwannomas development, other mechanisms likely also contribute to schwannoma development. For example, the constitutive activation of a variety other oncogenic signaling networks such as Ras-ERK, Rac1, src, β-catenin, and the mTOR protein kinase complex have been described in Merlin-deficient cells^20^. Additionally, reports in the literature implicate Merlin in establishing or maintaining cellular polarity^21^ and in regulating intracellular vesicular trafficking and the endocytic process, micropinocytosis^22,23^.

Our objective was to develop a more complete understanding of Merlin function in cell polarity, intracellular trafficking, and signal transduction. The challenge was in identifying those target proteins and pathways that are important for Merlin tumor suppressor function specifically when Merlin is active. We used a proximity biotinylation strategy coupled with a robust Merlin binding screening system to identify proteins that are proximal to both Merlin isoform 1 and isoform 2 at confluence, in contact inhibited cells when Merlin is active, but not in sub-confluent, growing cells when Merlin is inactive. We identified many novel Merlin proximal proteins that confirmed Merlin involvement in specific cell-cell and cell-substrate junctions, expanded the number of signal transduction pathways that Merlin may influence and highlight a critical role for Merlin in intracellular transport. We identified the small GTPases RalA and RalB as high affinity PIP_2_ dependent Merlin binding proteins and demonstrate that Merlin competitively inhibits RalB binding to its exocyst effectors Sec5 and Exo84. Merlin loss increased Ral activity in confluent Schwann cells and regulates the kinetics of exocytosis in a RalB dependent manner. Our results identify RalB as a novel, critical binding partner for active Merlin and raise the possibility that the RalA/B pathway may be targeted therapeutically to treat NF2-related schwannomatosis.

## Results

### Contact Inhibition Parameters

We defined the cell densities at which Merlin is active in the immortalized human Schwann cell line iHSC-1λ^24^ by comparing the growth parameters of Merlin knockout and scrambled gRNA control cell lines (Supplemental Figure 1). YAP1 staining confirmed a significant increase in the proportion of cytosolic YAP1 in control cells 48 hours after seeding at high density (10-20,000 cells per cm^2^) relative to low cell density (2,500 to 5000 cells per cm^2^ that was not apparent in the Merlin knockout cells (Figure 1A, B, Supplemental Fig. 2). We generated iHSC-1λ cell lines expressing Merlin isoform1 or isoform 2 with a recombinant ascorbate peroxidase (APEX2)^25^ fused to their C-terminus (Figure 1A). We measured APEX activity in lysates from these cell lines. Mer1-APEX and Mer2-APEX activity was roughly equivalent and four-fold less than control cell lysates expressing APEX alone (Figure 1B). Merlin iso1 and iso2-APEX proteins were expressed ∼1.6 and 2.0-fold over endogenous levels, as measured by densitometry (Figure 1C). There was no significant difference in the subcellular localization of endogenous Merlin (Figures 1D., E.), Merlin isoform 1-APEX (Figures 1F., G.) and Merlin isoform 2-APEX (Figures 1H., I.), ruling out artifacts caused by gross differences in subcellular localization.

**Figure 1.**
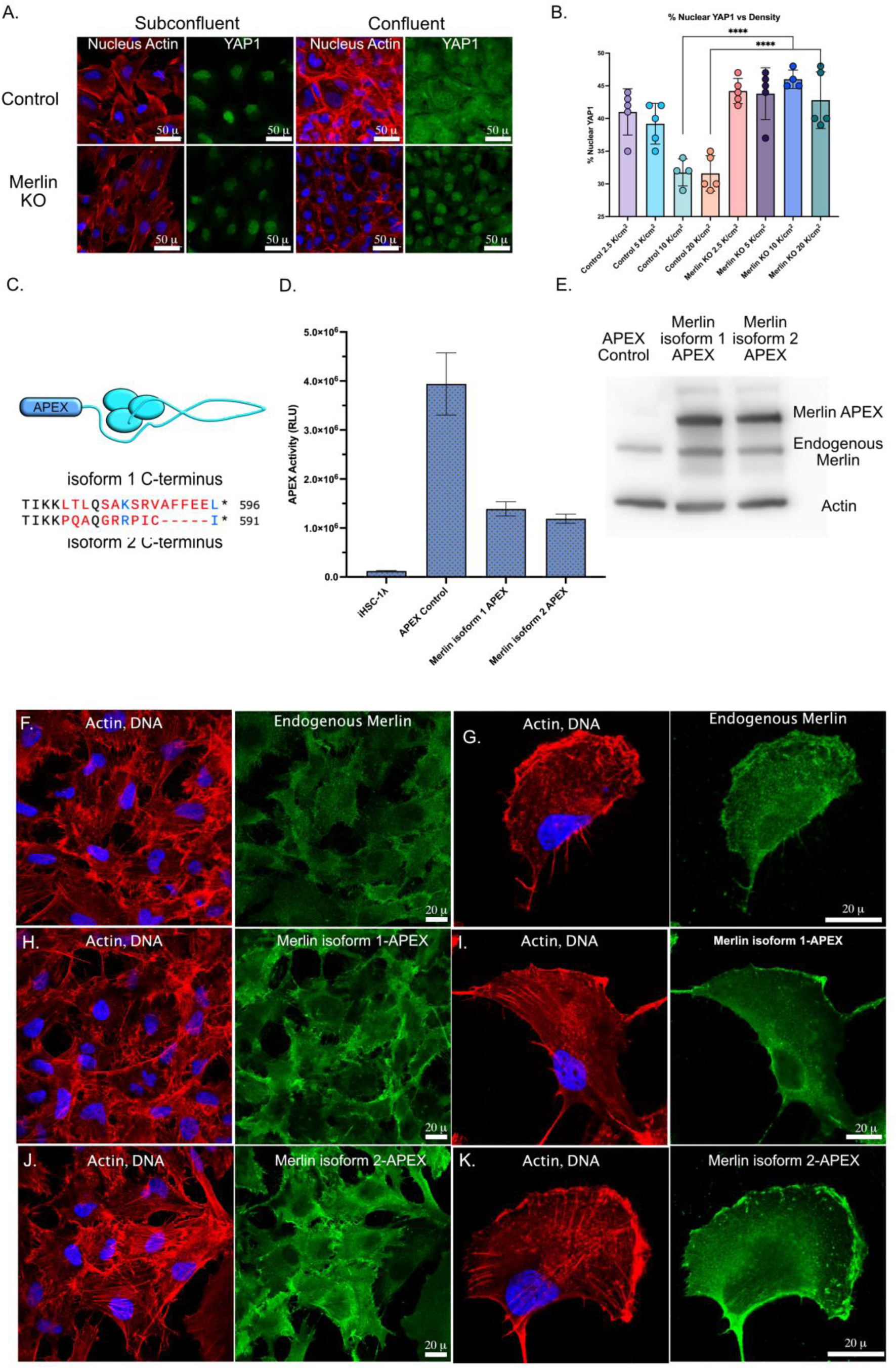
Merlin-APEX Cell Lines. A. Control and Merlin knockout cells plated at low cell density (2,500 cells per cm^2^) and high cell density (20,000 cells per cm^2^), incubated for 48 hours then fixed and stained with anti-YAP1 antibodies and counterstained for nuclei and actin with DAPI and phalloidin and confocal images were acquired with 60x objective. Scale bar = 50 µ. B. Quantitation of the nuclear to cytoplasmic ratio of YAP1 at different cell densities. Control and Merlin knockout cells were plated at 20,000, 10,000, 5,000 and 2,500 cells per cm^2^, incubated for 48 hours, stained with antibodies to YAP1 then confocal images were acquired with 20x objective, 4 fields per point. Acquisition parameters were constant for all images. To assess the proportion of YAP1 in the nucleus, the DAPI stained images were used to make a binary mask that was applied to the YAP1 image to measure the integrated density of the YAP1 signal specific to the nucleus. The proportion of nuclear staining was expressed as: % nuclear YAP1 = nuclear signal ÷ (total signal−nuclear signal). C. A schematic diagram representing Merlin with the proximity biotinylation enzyme APEX2 fused to its C-terminus. Below: the amino acid sequences for the Merlin isoform 1 and isoform 2 C-terminus. D. Normalized APEX activity in APEX control, Merlin isoform 1-APEX and Merlin isoform 2-APEX E. Immunoblot analysis of cell lysate from parental iHSC-1λ, and Merlin isoform 1-APEX and Merlin isoform 2-APEX expressing cells probed with antibodies to Merlin and β-actin as a loading control. F-K Subcellular location of Merlin in confluent monolayers (F., H., J.) and individual cells (G., I., K.) for endogenous Merlin (F., G.), Merlin isoform 1-APEX (H., I.) and Merlin isoform 2-APEX (J., K.).

### Proximity Biotinylation

iHSC-1λ cells expressing either Merlin iso1-APEX or Merlin iso2-APEX were plated at either 5,000 cells per cm^2^ or 15,000 cells per cm^2^ and incubated for 48 hours to produce either a sub-confluent, growing cell population or a confluent, contact inhibited cell population (Figure 2A). We initiated a 1-minute APEX mediated biotinylation reaction, then harvested cell lysates. Biotinylated proteins were purified by streptavidin affinity chromatography, washed extensively then processed for MALDI-TOF mass spectroscopy. These experiments were performed in triplicate. Western blots of aliquots showed significant biotinylated proteins with a range of molecular weights for both sub-confluent and confluent cells, with a noticeable increase in the confluent samples (Figure 2B).

**Figure 2.**
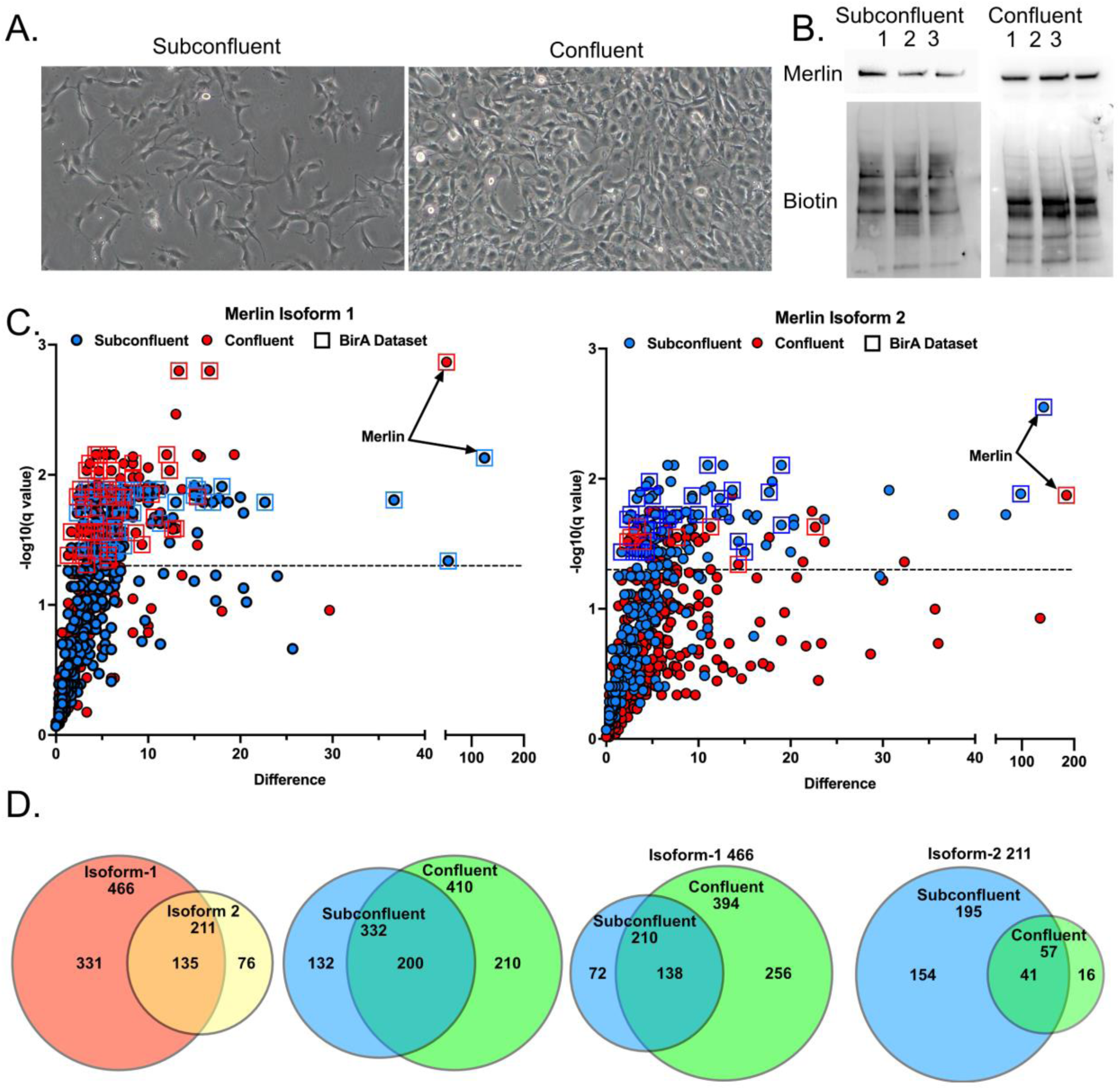
Merlin Proximity Biotinylation. A. Typical subconfluent (left) and confluent (right) cell densities for Merlin isoform 1-APEX proximity biotinylation experiments. B. Western blots of biotinylated proteins from Merlin isoform 1-APEX from subconfluent and confluent cells probed with antibodies to Merlin (top) and streptavidin to visualize biotinylated proteins (bottom). C. Merlin isoform 1 proximal proteins, displayed as Volcano plots showing statistical significance plotted against the peptide difference between the Merlin and control. The Y-axis plots the Q value (Q= − log_10_ *p*). The dotted line indicates the statistical significance threshold (p=0.05, Q=1.30). The X-axis plots the mean difference of normalized peptide counts per identified protein between Merlin-APEX expressing cells and APEX expressing control cells. Data from subconfluent cells is plotted in blue and from confluent cells is plotted in red. Proteins that had been identified in our published, BirA-based proximity biotinylation experiments are indicated by a surrounding square D. Venn diagrams depicting the number of proteins biotinylated by Merlin isoform 1-APEX (left graph) and Merlin isoform 2-APEX (right graph) greater than APEX only control (5% FDR) in confluent and subconfluent cells. The Venn diagrams show the number of unique and common proteins comparing (left to right) isoform 1 vs isoform 2, confluent vs subconfluent for all identified proteins, confluent vs subconfluent for isoform 1 and confluent vs subconfluent for isoform 2.

Merlin proximal proteins were identified from the mass spectroscopy data by the normalized total number of peptides mapping to each protein in the Merlin-expressing samples that were significantly greater than in the APEX control samples, as judged by T-test (p-value of <0.05 with a 5% false discovery rate, Supplemental Spreadsheet 1). The mass spectroscopy data for isoform 1 and isoform 2 proximal proteins is displayed in Volcano plots showing the mean normalized peptide difference between Merlin samples and APEX only control and the statistical significance (Q= − log_10_ *p*) (Figure 2C). We identified a total of 542 proteins across the four experimental samples that met the selection criteria. Of these, 81 proteins were also identified in our published proximity biotinylation dataset (indicated by boxed datapoints in Figure 2C)^14^. Furthermore, we identified multiple proteins reported to bind Merlin in the literature, including angiomotin^26^, Erbin^27^, YAP1^28^, TP53BP2^14^, PPP1R12A (Mypt1)^29^, Ezrin^30^ and Moesin^10^ (Table I), validating the new dataset. There were 466 proteins identified by isoform 1 and 410 by isoform 2 (Figure 2D). A total of 322 proteins were present in sub-confluent cells and 410 in confluent cells, with 200 proteins that overlapped. The number of proteins proximal to Merlin isoform 1 significantly increased in confluent cells, from 211 to 394 with 138 proteins identified at both cell densities but 256 identified exclusively in confluent cells. In contrast, 195 of the 210 proteins identified by Merlin isoform 2 were in sub-confluent cells, with only 57 in confluent cells. This suggests that the isoforms might have different functions in addition to their shared tumor suppressor function, and that isoform 1 may be more active in confluent cells while isoform 2 is more active in sub-confluent cells.

**Table I.** Gene Ontogeny Categories. Merlin proximal proteins mapped to each gene ontogeny term, the specific GO databases, including their description, specific accession number and the p-value associated with the analysis.

| Network | Hub | Database | Acession | Proteins | FDR value |
| --- | --- | --- | --- | --- | --- |
| Cell Junctions | Cell Junction | GO Cellular Component | GO:0030054 | 206 | 1.83E-57 |
| Cell Junctions | Adherens Junction | GO Cellular Component | GO:0005912 | 40 | 1.85E-20 |
| Cell Junctions | Focal Adhesion | KEGG Pathways | hsa04510 | 30 | 2.16E-11 |
| Cell Junctions | Tight Junction | KEGG Pathways | hsa04530 | 28 | 6.78E-12 |
| Trafficking | Intracellular Transport | GO Biological Process | GO:0046907 | 87 | 1.35E-10 |
| Trafficking | Membrane Trafficking | Reactome Pathways | HSA-199991 | 80 | 5.53E-26 |
| Trafficking | Endocytosis | GO Biological Process | GO:0006897 | 44 | 3.87E-10 |
| Trafficking | Clathrin-Mediated Endocytosis | Reactome Pathways | HSA-8856828 | 28 | 2.70E-12 |
| Trafficking | Exocytosis | GO Biological Process | GO:0006887 | 26 | 6.85E-07 |
| Trafficking | Vesicle Docking Involved in Exocytosis | GO Biological Process | GO:0006904 | 9 | 4.40E-04 |
| Signaling | Signal Transduction | Reactome Pathways | HSA-162582 | 175 | 4.02E-28 |
| Signaling | Kinase | GO Molecular Function | GO:0016301 | 54 | 2.53E-07 |
| Signaling | Kinase | UniProt Keywords | KW-0418 | 42 | 9.71E-06 |
| Signaling | Transmembrane RTK Signaling | GO Biological Process | GO:0007169 | 40 | 1.05E-08 |
| Signaling | Molecular Adaptor Activity | GO Molecular Function | GO:0060090 | 36 | 9.42E-08 |
| Signaling | Phosphatidylinositol Binding | GO Molecular Function | GO:0035091 | 25 | 3.51E-05 |
| Signaling | HIPPO | WikiPathways | WP4541 | 23 | 6.10E-10 |
| Signaling | PI3K-Akt signaling Pathway | KEGG Pathways | hsa04151 | 23 | 1.20E-03 |
| Signaling | Small GTPase | InterPro Domains | IPR001806 | 20 | 5.69E-06 |
| Signaling | HIPPO | KEGG Pathways | hsa04390 | 18 | 1.25E-05 |
| Signaling | HIPPO | WikiPathways | WP4540 | 17 | 9.26E-07 |
| Signaling | HIPPO | Reactome Pathways | HSA-2028269 | 7 | 1.10E-04 |

### Functional Enrichment Analysis

To begin to understand the potential functional significance of these proteins we performed a STRING based functional enrichment analysis on the Merlin proximal proteins^31^, displayed as a proximity network^32^ (Figure 3A). The proximal proteins were color coded for gene ontogeny terms to indicate proteins functioning in cell junctional complexes, intracellular trafficking and signal transduction (Table I). We identified 206 proteins associated with cell junctions, including 40 adherens junction proteins, 30 focal adhesions proteins and 28 tight junction proteins (Table I). These included N-cadherin, catenin-α, catenin-β, catenin-δ, integrin-β1, integrin-α2 and the ephrin receptors EPHA2, EPHB2 and EPHB4 (Table II, Supplemental Spreadsheet 1). We also identified 137 proteins involved in intracellular transport or membrane trafficking, (87 and 80 proteins respectively), and 107 proteins more specifically mapping to endocytosis or exocytosis (Table I). These included 10 members of the Rab family of small GTPases, most prominently the endocytic Rab7a and the recycling/exocytic Ral11b (Table II, Supplemental Spreadsheet 1). Other trafficking proteins include the AP-2 complex subunits AP2A1 and AP2A2 and proteins critical for exocytosis, including 3 out of the 8 components of the exocyst, EXOC2, EXOC4 and EXOC7 (Table II, Supplemental Spreadsheet 1). Significantly, we identified 173 signal transduction proteins (Table I), with 48 mapping to the HIPPO pathway, including angionmotin, ASPP2, and YAP1 (Table II, Supplemental Spreadsheet 1). Surprisingly, neither of the HIPPO core kinases, Lats1/2 or Mst1/2, were identified. There were 40 members of the receptor tyrosine kinase pathway and 25 members of the PI3K/Akt3 pathways (Table I); a total of 54 proteins were kinases or had kinase activity. These included growth factor receptors (EGFR, MET, and PDGFRβ), and protein kinases (PAK2, JAK1, integrin linked kinase, ILK1 and myosin light chain kinase MYLK) (Table II, Supplemental Spreadsheet 1). There were 25 phosphatidylinositol binding proteins and 36 molecular adapters (Table I). There were also 19 small GTPases, including 10 Rab family members plus CDC42, NRAS, RHOA, RRAS2, RAN, RAP1A, RAP2A, RALA and RALB (Table II, Supplemental Spreadsheet 1).

**Figure 3.**
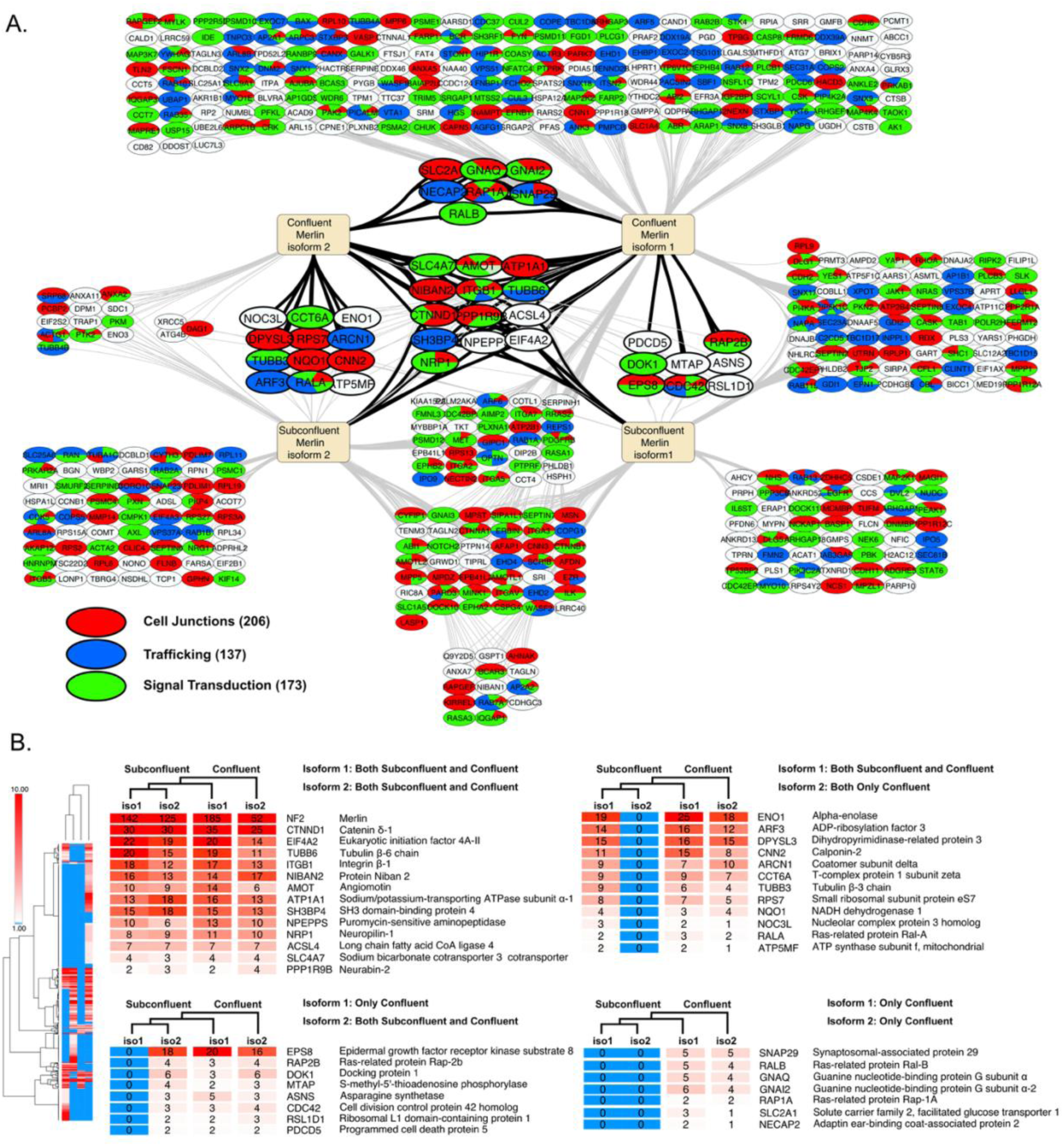
Merlin Proximity Network. A. Merlin proximal proteins displayed as a proximity network for Merlin isoform 1 (right) and isoform 2 (left) in subconfluent (bottom) and confluent (top) cells. The nodes are color coded based on gene ontogeny terms derived from STRING based functional enrichment analysis for cell junctional proteins (red), signal transduction proteins (green) and intracellular transport (blue). Proteins sets that are proximal to both isoform 1 and isoform 2 in confluent cells are highlighted. B. Hierarchical cluster analysis showing four subsets of proteins proximal to both isoform 1 and isoform 2 in confluent cells are shown. The mean normalized peptide numbers are displayed within the box the relevant gene symbol and protein name are presented. The blue cells indicate no significant difference with APEX alone controls.

**Table II.**
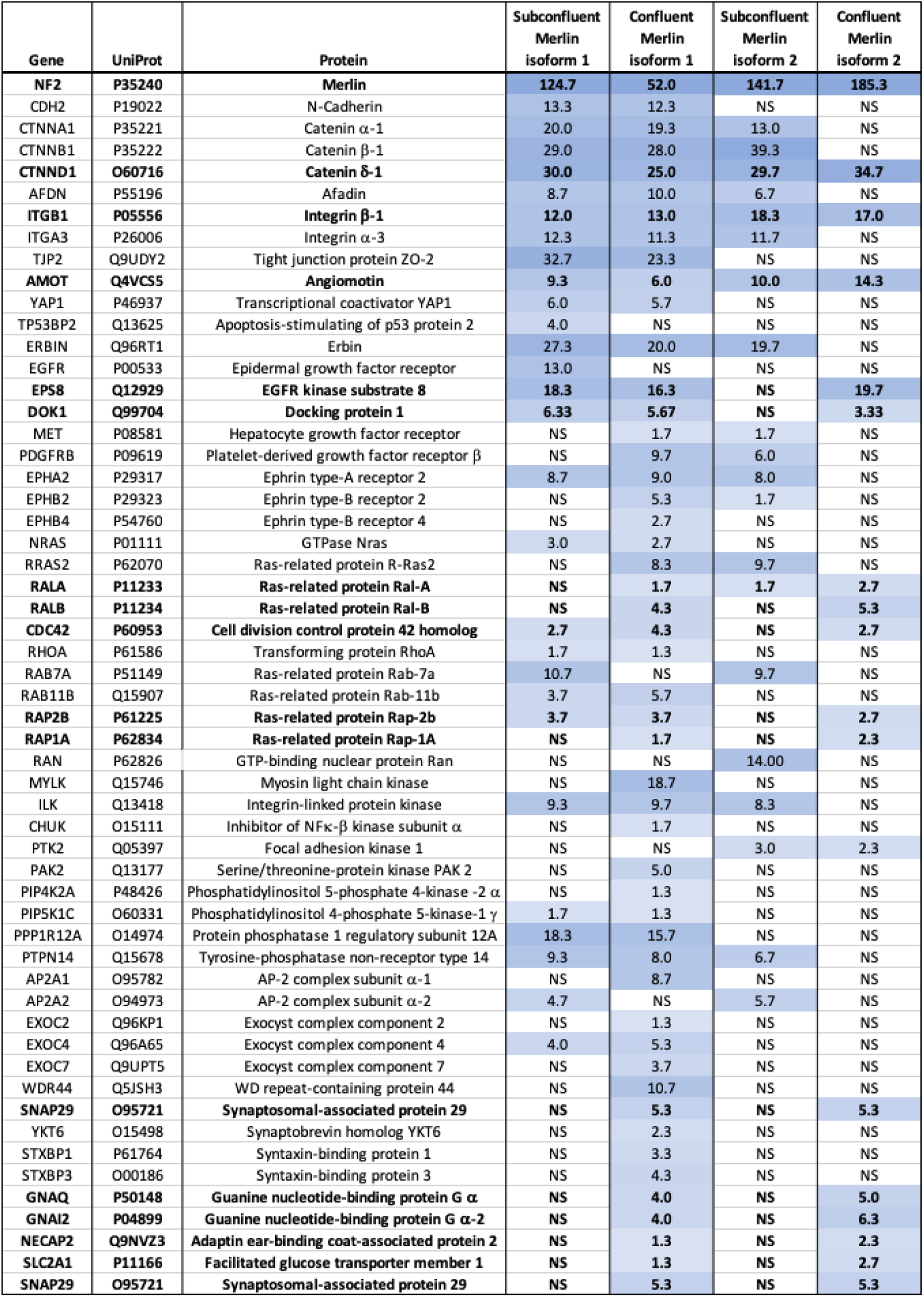
Selected Merlin Proximal Proteins. Merlin proximal proteins that were mentioned in the text, showing the mean normalized peptide count per 10,000 in each experimental condition, the intensity of the blue background is proportional the peptide count. NS indicates no significant difference in mean peptide number relative to the APEX only control. Proteins highlighted in bold are proximal to both isoform 1 and isoform 2 in confluent cells.

A subset of this network consisted of 41 proteins biotinylated by both isoform 1 and isoform 2 in confluent cells. We hypothesized that this cluster would contain the proteins most likely to interact with active Merlin. Hierarchical cluster analysis revealed four subgroups (Figure 3B). There were 14 proximal to both isoform 1 and isoform 2 in sub-confluent and confluent cells, including the Merlin binding protein angiomotin and upstream cell junctional complexes like integrin-β1 and catenin-δ (Figure 3B, Table II). Two sets totaling 20 proteins were proximal to both isoform 1 and isoform 2 in confluent cells and either isoform 1 or isoform 2 in sub-confluent cells. These included the molecular adapter proteins DOK1 and EPS8, the calcium binding protein calponin-2 and the small GTPases cdc42, Rap2 and RalA (Figure 3B, Table II). Finally, there was a subset of 7 proteins are proximal to both isoform 1 and isoform 2 exclusively in confluent cells. These included the small GTPases RalB and Rap1a, the v-SNARE protein SNAP-29, the large GPCR subunits GNAQ and GNAI2, the facilitated glucose transporter SLC2A1 and the endocytosis associated protein NECAP2 (Figure 3B, Table II).

### Merlin Binding

We hypothesized that Merlin proximal proteins that are critical to its tumor suppressor function are likely to be signaling proteins that physically bind to active Merlin. We therefore focused screening experiments on a selection of Merlin-proximal cell junctional complex proteins, ERM proteins, kinases, phosphatases and small GTPases. Merlin is activated by PIP_2_ binding^15^ which induces a conformational change^16^ that allows dimerization^11^ and increases its affinity for binding^33^. Therefore, we screened for direct binding to Merlin binding using purified proteins in the presence and absence of PIP_2_^11,14^(Figure 4A-C). Figure 4D presents the results of 35 of these assays performed in triplicate, expressed relative to the GFP negative control. We evaluated known Merlin binding proteins angiomotin, Lats1, ASPP2 and YAP1. Merlin consistently showed the highest relative affinity for angiomotin, > 65-fold above control, in a PIP_2_ independent manner. Merlin also showed relatively high affinity for Lats1, but significantly less for ASPP2 and YAP1. There was a marked PIP_2_-dependent increase in affinity for YAP1 (Figure 4D). In contrast, the cell adhesion complex proteins N-cadherin, β1-integrin, catenin-α, catenin-β and catenin-δ showed low relative affinity binding, (2-4-fold) and minimal PIP_2_ effect. Similarly, Merlin showed low relative affinity binding and small increases in the presence of PIP_2_ to the ERM proteins Ezrin, Moesin, and the adapter protein Erbin. The same pattern was seen for kinases (PTK2, CHUK, ILK, MYLK, AKT1 and AKT2), and phosphatases (PP1A and MYPT) tested. All these proteins bound Merlin greater than they bound control, but with relatively low affinity and little increase in the presence of PIP_2_.

**Figure 4.**
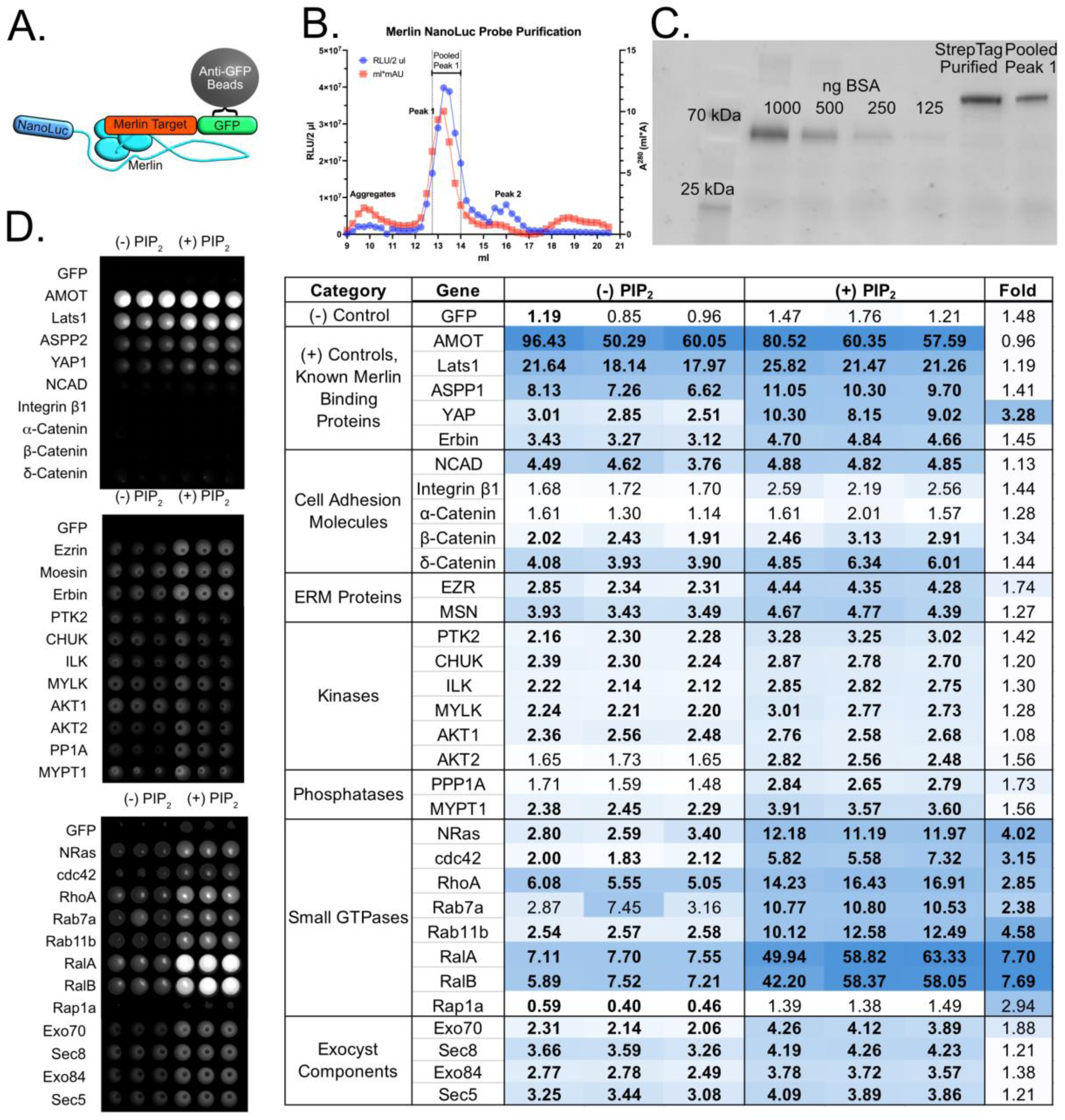
Direct Merlin Binding Assays. A. A schematic diagram of the direct Merlin binding assay showing the Merlin-NanoLuc probe, the GFP fused bait protein bound to anti-GFP nanobody bound beads. B. Merlin-NanoLuc probe purification by gel filtration showing the fractionation of affinity purified Merlin-NanoLuc activity and A^280^, showing the pooled peak 1 used in the binding assays. C. Aliquots of purified Merlin-NanoLuc proteins stained for total protein. Left to right: Marker, BSA titration from 1000 to 125 ng, Merlin-NanoLuc from the initial affinity purification step, and Merlin-NanoLuc pooled from peak 1. D. Luminescence images from 96 well plates containing triplicate Merlin binding assays in the presence and absence of di8-PIP_2_ for 33 putative Merlin binding proteins. E. A heat map showing luminometer data for Merlin binding activity in triplicate. Data are normalized to the mean of the GFP control for the reactions shown in D. The intensity of the blue color is proportional to the relative binding. The data highlighted in bold indicates statistical significance (T-test, p <= 0.05). The last column shows the fold increased binding in the presence of PIP_2_, bold numerals are statistically significant.

In contrast, the small GTPases, N-Ras, cdc42, RhoA, Rab7a, Rab11b, RalA and RalB showed significant PIP_2_-dependent binding. Most striking was the high relative affinity and the significant increase in binding in the presence of PIP_2_ that Merlin showed for RalA and RalB. Given the strong binding between Merlin and RalA and RalB, we searched for Ral-GAP, -GEF and effector proteins in our Merlin-proximal dataset. We identified Exo70, Sec8 and the Ral effector Sec5, three components of exocyst complex (Table II, Supplemental Spreadsheet 1). The exocyst is a conserved 8 protein complex that mediates the initial tethering of the vesicle to the plasma membrane^34^, stimulates SNARE complex assembly^35^ and functions in both constitutive^36^ and regulated exocytosis^37^. Merlin binding to Exo70, Sec8 and to the Ral effectors Sec5 and Exo84 was detectable and showed a slight increase in the presence of PIP_2_. We conclude that RalA and RalB are good candidates for critical signal transduction proteins that interact directly with active, PIP_2_-bound Merlin.

### Merlin Ral Interactions

We took advantage of the NanoLuc-GFP binding assay to perform bioluminescence resonance energy transfer (BRET) to confirm close interaction between Merlin-NanoLuc and GFP RalA or GFP-RalB in complex. There was significant excitation of the GFP fluorescence, peaking at 510 nm in both GFP-RalA and GFP-RalB proteins when in complex with Merlin-NanoLuc, indicating that Merlin forms close complexes with both RalA and RalB (Figure 5A). Given the relatively high affinity that Merlin has for RalA and RalB we tested whether Merlin affects Ral activity. We assayed for both RalA and RalB activity in control and Merlin knockout iHSC-1λ cells, at either sub-confluent or confluent cell densities. In sub-confluent cells, both RalA-GTP and RalB-GTP levels was equal between Merlin knockout and control cells, but in confluent cells RalA/B-GTP levels were significantly increased in Merlin knockout cells relative to control cells (Figure 5B). Increased RalA activity is consistent with reports in the literature^38^. Since our proximity biotinylation data identified RalB as proximal to both Merlin isoforms only confluent cells (Figure 3), we chose to focus on RalB going forward.

**Figure 5.**
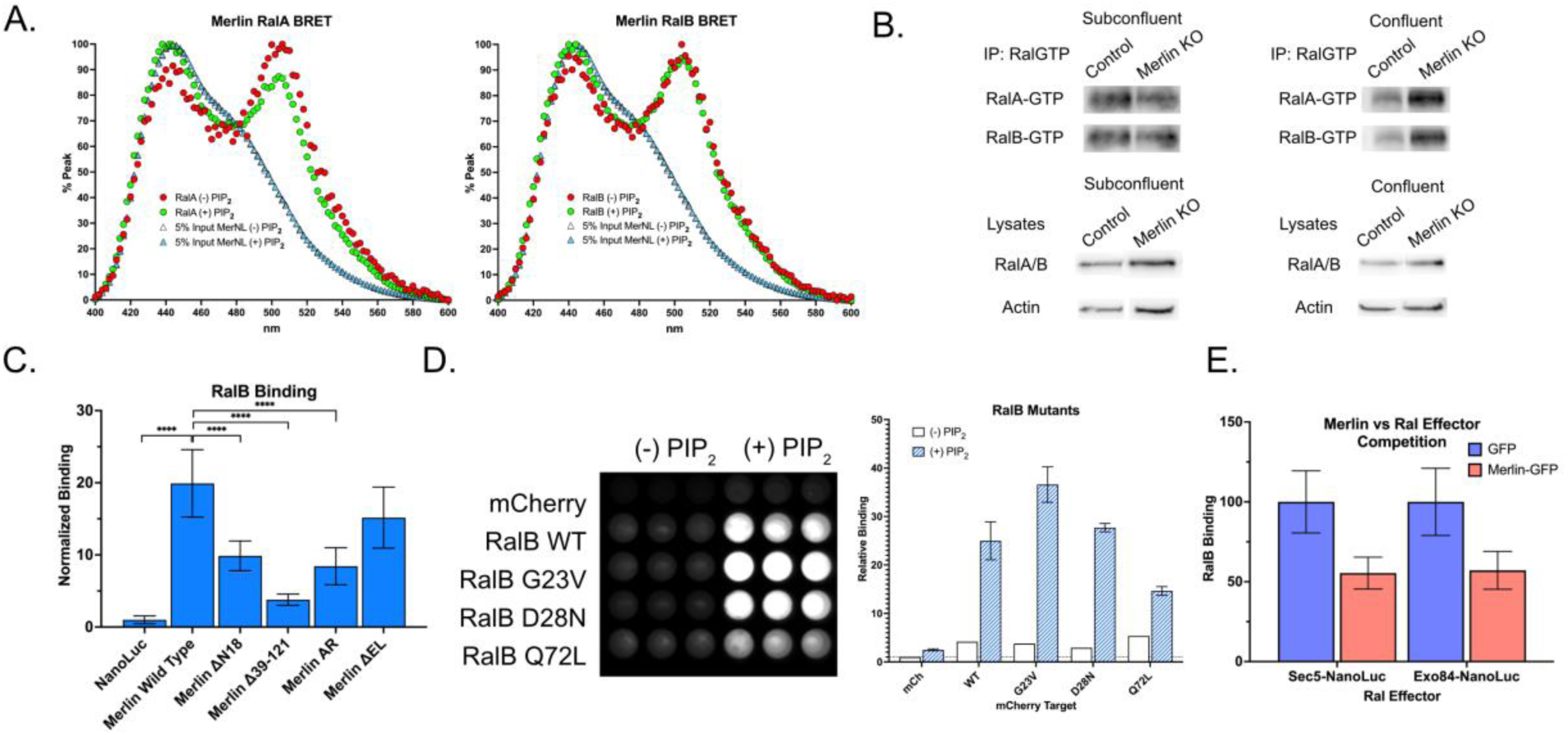
Merlin-RalA/B Interaction. A. BRET assays of Merlin-NanoLuc:GFP-RalA (left) and Merlin-NanoLuc:GFP-RalB (right) showing excitation of the GFP-Ral proteins by the luminescence of Merlin-NanoLuc in complex. B. RalA and RalB activity in Control and Merlin KO cells in subconfluent and confluent conditions. C. Co-immunoprecipitation of NanoLuc-RalB with wild type and mutant Merlin-GFPs. Left to right: NanolLuc alone control, wild type Merlin, an N-terminal deletion mutant Δ18, the FERM domain deletion mutant Δ39-121, the closed conformation mutant AR and the open conformation mutant ΔEL. D. Merlin binding for RalB wild type, the G23V and Q72L active mutants and the D28N dominant negative mutant showing an image of the luminiescence (left) and normalized Merlin binding activity (right) in the presence and absence of di8-PIP_2_. E. Co-immunoprecipitation of mCherry-RalB with the exocyst effectors Sec5-NanoLuc and Exo84-NanoLuc in the presence of either control GFP of Merlin-GFP.

The Merlin-RalB interaction was impaired by deletion of Merlin’s unique 20 N-terminus (Merlin-ΔN18, Figure 5C) and lost in an in-frame deletion within the FERM domain (Merlin-Δ39-121, Figure 5B), suggesting that RalB binds to the Merlin FERM domain and that the Merlin N-terminus is necessary for this interaction. RalB binding was also impaired to the closed conformation mutant Merlin-AR^8^ (Figure 5C). In contrast, the open conformation mutant Merlin-ΔEL^8^ was unaffected (Figure 5C). These results suggest that RalB preferentially binds to Merlin’s open conformation and are consistent with the PIP_2_-dependent nature of this interaction.

RalA and RalB are small G-proteins, which cycle between a GTP-bound active form and a GDP-bound inactive form. To determine if Merlin specifically binds to active or inactive RalB we performed Merlin-binding assays using 2 constitutively active mutants (RalB^G23V^ and RalB^Q72L^) and a dominant negative mutant (RalB^D28N^). All 3 RalB proteins bound to Merlin in the presence of PIP_2_, with slight differences in relative binding efficiencies but no correlation to RalB activation status (Figure 5D). These results suggest that Merlin binds RalB in the presence of PIP_2_, suppresses RalB activity during contact inhibition of growth, and that in the absence of Merlin this negative regulation is lost. Next, to determine if Merlin binding either facilitates or inhibits RalB interaction with its exocyst effector proteins, we performed binding assays between RalB and Sec5 or Exo84 in the presence or absence of Merlin. These experiments showed a significant, 50%, reduction of RalB binding to Sec5 and Exo84 in the presence of Merlin (Figure 5E). This suggests that Merlin may function to competitively inhibit RalB binding to its exocyst effectors, Sec5 and Exo84.

To determine if endogenous Merlin co-localizes with the Ral GTPases, iHSC-1λ cells were co-stained with antibodies that detect Merlin and pan-RalA/B. Merlin was distributed on the ventral surface of the plasma membrane in a distinct punctate pattern and co-localized, in part, to cell junctions that can contain N-cadherin (Figure 6A). The exocyst marker Exo70 was also localized to the ventral plasma membrane, where it showed a punctate pattern with significant co-localization with endogenous Merlin (Figure 6B). Antibodies detecting RalA/B showed a similar ventral punctate staining pattern, and significant overlap with Merlin staining (Figure 6C). To better verify overlap, we acquired super-resolution microscopic images of Merlin and Ral A/B staining. This confirmed significant co-localization between Merlin and RalA/B, co-localized in ventral puncta (Figure 6D). This subcellular localization data is consistent with Merlin-Ral interaction and a role for Merlin and RalA/B in the process of exocytosis.

**Figure 6.**
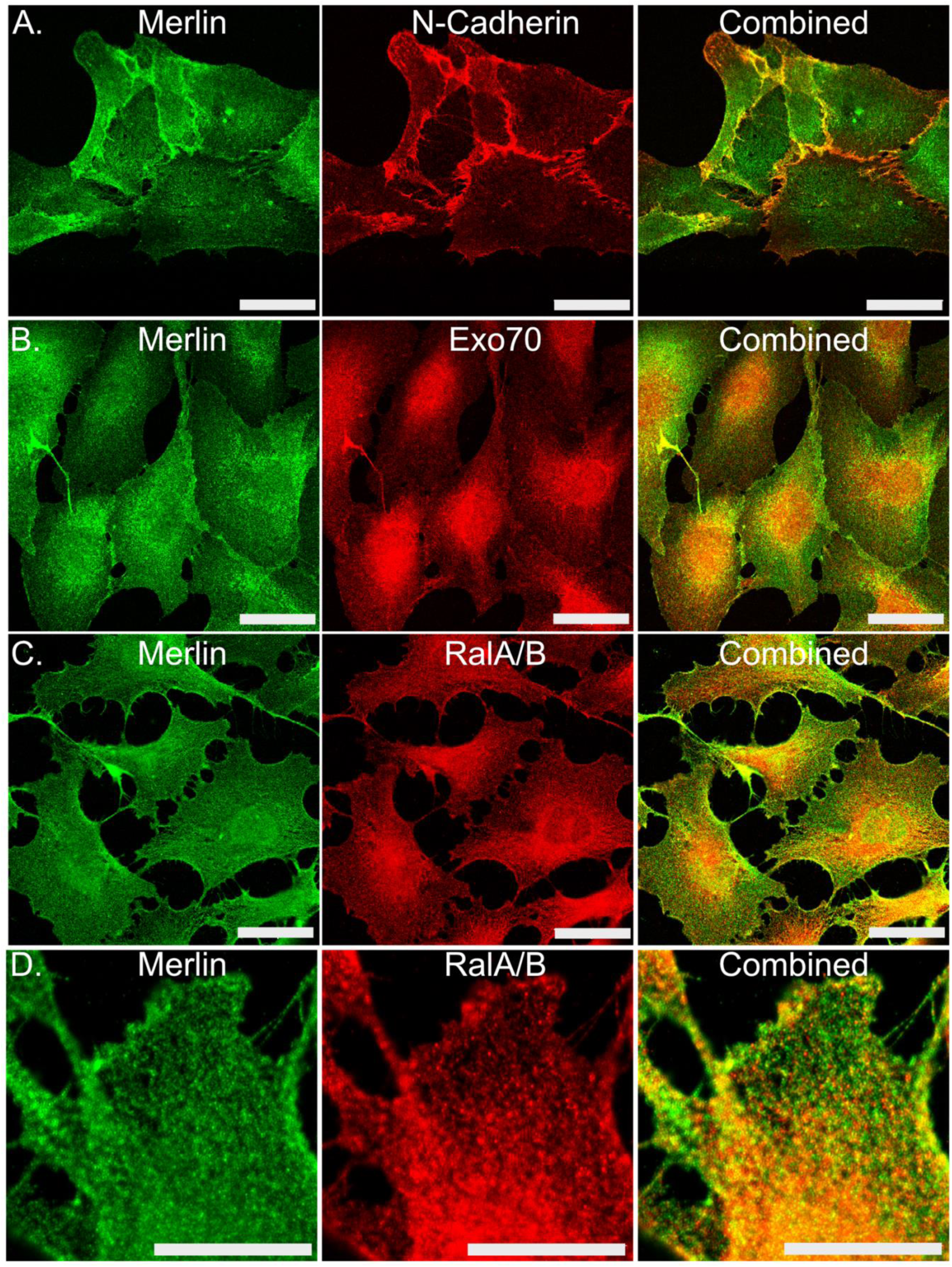
Subcellular localization of Endogenous Merlin, N-cadherin, Exo70 and RalA/B. A. Confocal images showing the pattern of co-localization of endogenous Merlin (green) and N-Cadherin (red) in the immortalized human Schwann cell line, iHSC-1λ. Bar = 50 µ. B. Confocal images showing the pattern of co-localization of endogenous Merlin (green) and Exo70 (red) in the immortalized human Schwann cell line, iHSC-1λ. Bar = 50 µ. C. Confocal images showing the pattern of co-localization of endogenous Merlin (green) and RalA/B (red) in the immortalized human Schwann cell line, iHSC-1λ. Bar = 50 µ. D. NSPARC super-resolution showing the co-localization of Merlin (green) and RalA/B (red) in iHSC-1λ. Bar = 20 µ

### Merlin Regulates Exocytosis via RalB

The ability of Merlin to bind to RalB in a PIP_2_-dependent manner, to competitively inhibit RalB binding to its exocyst effectors Sec5 and Exo84 and the co-localization of endogenous Merlin with RalA/B punctate structures on the ventral plasma membrane strongly suggested that Merlin might regulate exocytosis via RalB at the exocyst. To test this idea, we used a vesicular membrane marker, VAMP2, fused to the pH sensitive GFP mutant, pHluorin, to measure exocytosis^39^. Using this marker, endocytic events are indicated by transient increases in fluorescence in small areas on the ventral cell surface, which can be visualized in live cells by TIRF microscopy over for 1-2 min., in images taken at 100 msec. intervals (Figure 7A). We compared exocytosis between control and Merlin knockout iHSC-1λ cells and tabulated the number, size, intensity and duration of events. Examples mapping exocytic events in control and Merlin knockout cells are presented in Figure 7B and C. There was no significant difference in the number, size or intensity of the exocytic events between control and Merlin knockout. However, we identified a significant difference in event duration. This result is apparent in a frequency histogram of these data (Figure 7D). In control cells ∼59% of the events lasted between 1000 and 2000 msec., ∼33% from 2000 to 3000 msec. and ∼8% lasted longer than 3000 msec. In Merlin knockout cells the figures are ∼39%, 41% and 20% respectively. The median duration for control and Merlin knockout is 1300 msec. vs 1600 msec with mean values of 1612 vs 1974 respectively (Figure 7E). These data show that loss of Merlin affects the kinetics of exocytosis.

**Figure 7.**
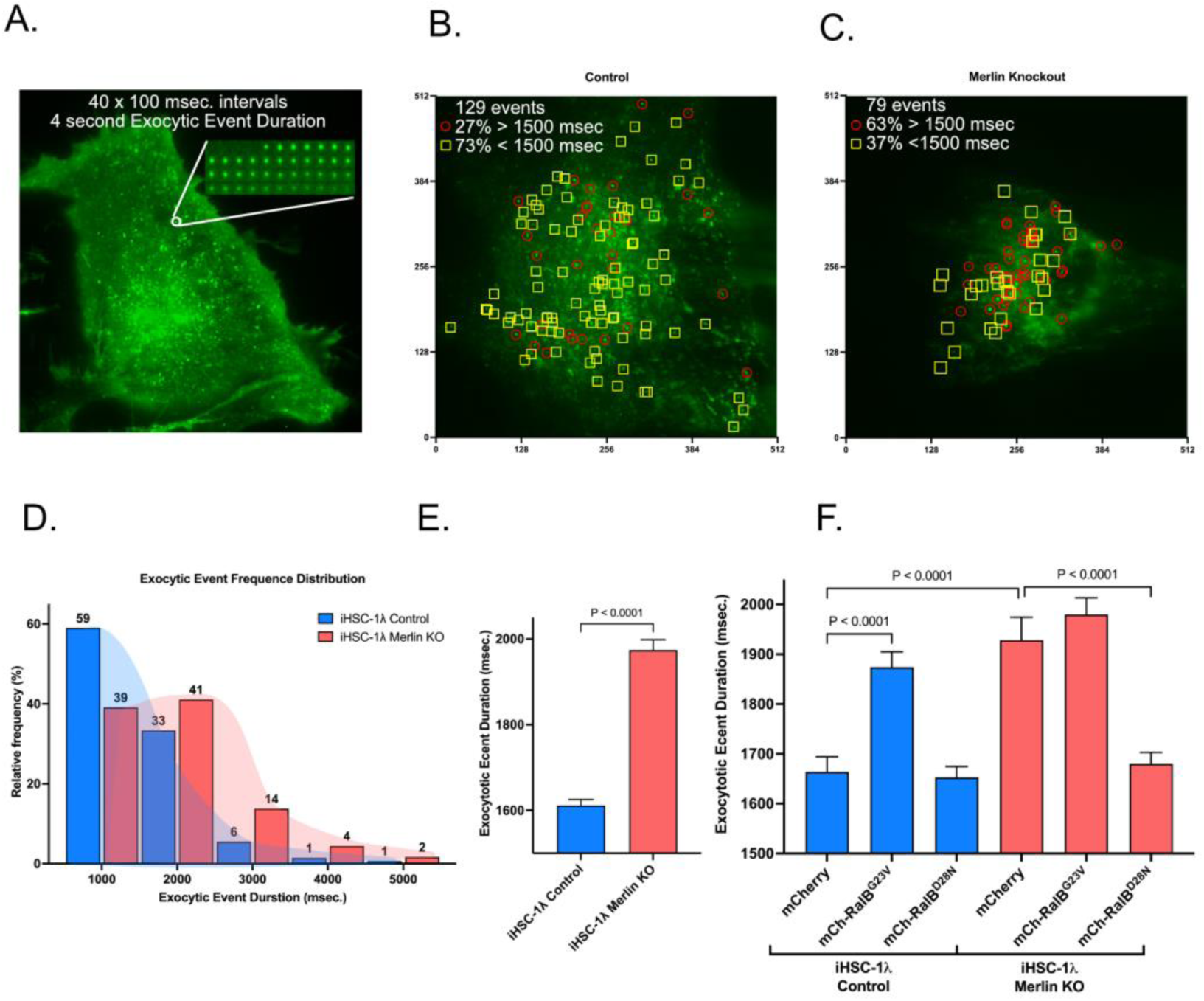
Merlin Regulation of Exocytosis Kinetics. A. An image of showing a summation of VAMP2-pHlourin exocytic events over a 1-minute time lapse at 100 msec intervals for a total of 601 frames per cell in a live iHSCL-1λ Schwann cell. Exocytic events were defined by the appearance of an increase in green fluorescence of at least 4-fold above background in a 4-8 square pixel in area for up to 11 seconds with a R^2^ of at least 0.8 for decay kinetics and 0.75 for Gaussian distribution using ExoJ plugin of Image J. Inset: time course of a single, 4 second, exocytic event shown at 100 msec intervals. B. An example of the total number of exocytic events identified in a control iHSC-1λ Schwann cell over a 1-minute time course at 100 msec. intervals. The red circles identify event durations greater than 1.5 sec. and the yellow squares identify event durations less than or equal to 1.5 sec. C. An example of the total number of exocytic events identified in a Merlin knockout iHSC-1λ Schwann cell over a 1-minute time course at 100 msec. intervals. The red circles identify event durations greater than 1.5 sec. and the yellow squares identify event durations less than or equal to 1.5 sec. D. Frequency histogram depicting the exocytic event frequency comparing iHSC-1λ control and iHSC-1λ Merlin knockout cells showing a shift to longer event duration events in Merlin knockout cells. E. Mean exocytic event duration plus SEM of iHSC-1λ control and iHSC-1λ Merlin knockout cells. F. Mean exocytic duration of iHSC-1λ control and iHSC-1λ Merlin knockout cells expressing control mCherry, mCherry-RalB, the active mutant mCherry-RalB^G23V^ and the dominant negative mutant mCherry-RalB^D28N^.

**Figure 8.**
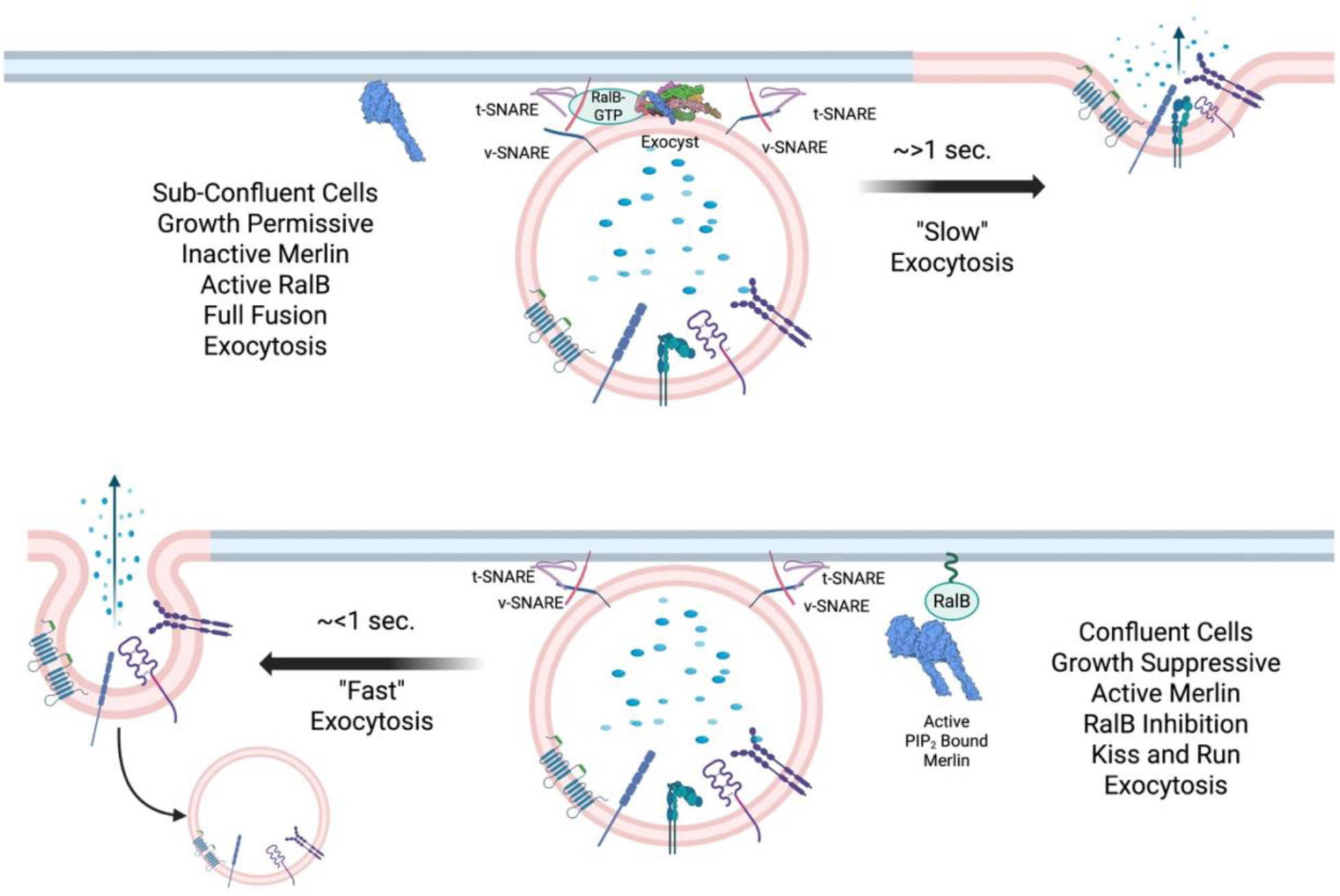
Merlin-Ral Regulation of Exocytosis. A. A schematic diagram of Merlin regulation of exocytosis by competitive inhibition of RalB. In actively growing, sub-confluent cells (top), Merlin is inactive and RalB mediated exocytosis leads to the export of integral membrane proteins and lipid plasma membrane components to the cell surface, supporting the expansion of the plasma membrane that is necessary for active cell growth. Contact-inhibited cells that are not actively growing (bottom) do not require plasma membrane expansion, Merlin is activated by PIP_2_ mediated dimerization, binds to RalB, to prevent association with the exocyst and inhibit full fusion exocytosis.

To test if RalB activity is necessary for this effect on the kinetics of exocytosis, we generated control and Merlin knockout cell lines co-expressing VAMP2-pHluorin with mCherry fused to active RalB^G23V^, dominant negative RalB^D28N^ or an mCherry control. The control cells expressing mCherry alone had a mean exocytic duration of 1664 msec (Figure 6H). Expression of active RalB^G23V^ significantly increased this value to 1874 msec., phenocopying the effect of Merlin loss (Figure 7F). In contrast, Merlin knockout cells expressing mCherry alone had a mean exocytic duration of 1928 msec and expression of active RalB^G23V^ did not significantly change this value (1979 msec). In contrast, Merlin knockout cells expressing dominant negative RalB^D28N^ had a mean exocytic event duration of 1608 msec, reverting to control event duration. We conclude that Merlin regulates exocytic kinetics by inhibiting RalB activity.

## Discussion

### Merlin Proximal Proteins

We identified 542 Merlin-proximal proteins in Schwann cells, including known Merlin proteins and proteins identified in a previous Merlin proximity biotinylation study^14^, validating the dataset. Overall, we confirmed and extended earlier data by identifying the cell surface receptors for cell to cell and cell to substrate cell junctions (Table I). A striking new result was the strong representation of Merlin-proximal proteins involved in intracellular vesicular trafficking, an observation that is consistent with data from the literature^22,23,40–42^. Finally, we identified 165 proteins involved in general signal transduction, the majority of which are kinases, molecular adapters or phosphatidylinositol binding proteins, including 48 proteins associated with the HIPPO pathway and 19 small GTPases, including 10 members of the Rab subfamily. The other small GTPases were, CDC42, NRAS, RRAS2, RALA, RALB, RAN, RAP1A, RAP2A, and RHOA. Notable by its absence was RAC1, the small GTPase that was most often associated with Merlin in the early literature^43^. The absence of a protein like Rac1 and a known Merlin binding protein like Lats1/2 does cannot exclude these proteins as Merlin interactors because of biases inherent in the proximity biotinylation system^44^.

### Merlin Binding Proteins

Our goal was to identify proteins that interact with Merlin when it is activated by PIP_2_-mediated dimerization^11^. Direct binding assays allowed us to test for interaction in the presence and absence of PIP_2_, that is, to test for binding to active Merlin^11^. Since PIP_2_ signaling is transient and spatially restricted to membranes, interactions that are increased when PIP_2_ is present are likely to represent a small but critical proportion of Merlin molecules within cells. Because the assay is quantitative and, although we did not formally measure K_d_, we were able to compare the relative affinity between Merlin and its binding proteins and to reveal a hierarchy of Merlin binding affinities. Anigomotin showed the highest affinity for Merlin. Binding was independent of PIP_2_, suggesting a distinct function for the Merlin-angiomotin complex. The HIPPO pathway proteins Lats1, ASPP2 and YAP1 showed strong Merlin binding, albeit with lower affinity than angiomotin; of these proteins only YAP1 showed significantly increased binding in the presence of PIP_2._

The small GTPases RalA and RalB had a higher relative affinity for Merlin, roughly 7-fold over the GFP control, which increased dramatically to more than 50-fold over control in the presence of PIP_2_. This level of binding is significantly higher than any other tested proximal proteins and on par with that of angiomotin. RalA and RalB are proximal to both isoform 1 and isoform 2 in confluent cells, with RalB being one of the seven proteins that are proximal to both isoforms exclusively in confluent cells. Endogenous Merlin colocalizes with RalA/B on the ventral surface of cells, in a punctate pattern similar to that of the exocyst component, Exo70, and consistent with a subcellular localization where exocytosis occurs. RalA/B activity is significantly increased in confluent Merlin knockout cells, but not in sub-confluent cells. Increased RalA activity upon Merlin loss was described more than 20 years ago^38^, although the mechanism described in that paper, Merlin binding to and inhibiting the Ral activator, RalGDS, is not supported by our data since RalGDS is not among the Merlin proximal proteins we identified. However, we cannot rule this out as a mechanism by which RalA/B is activated in confluent cells upon Merlin loss. Merlin competitively inhibited RalB binding to the exocyst effectors Sec5 and Exo84. Since both RalA and RalB bind to Sec5 and Exo84 it is likely that Merlin competitively inhibits RalA binding to the exocyst as well. Loss of Merlin caused a RalB dependent change in exocytosis kinetics that is phenocopied by active RalB in wild type cells. Taken together these data identify RalA and RalB as a key binding partner for active Merlin and show that Merlin regulates exocytosis in a RalB dependent manner (Figure 3B).

### RalA and RalB

Both RalA and RalB play critical roles a diverse set of cellular functions including proliferation, cell survival, differentiation motility and cellular polarization^45^. Mechanistically, many of these biological effects are dependent upon RalA/B regulation of endocytosis^46^ and exocytosis^47^. Ral regulation of endocytosis is achieved via the Ral effector RalBP which binds to the endocytic machinery via AP-2 to mediate EGF receptor mediated endocytosis^48^. RalBP was not among the Merlin proximal proteins we identified. Rather, the three exocyst components, Exo70, Sec8 and the Ral effector protein Sec5 were identified as Merlin proximal. These exocyst components plus the other Ral effector, Exo84, are all low affinity Merlin binding proteins (Figure 4), suggesting that the Merlin-Ral interaction may affect exocytosis rather than endocytosis, although the close coordination between endocytosis and exocytosis suggests that we cannot exclude a role for Merlin-Ral interaction in endocytosis as well as exocytosis.

The exocyst is responsible for tethering the exocytic vesicle to the inner face of the plasma membrane. It is a critical point of regulation via its interactions with vesicular and plasma membrane proteins, phosphoinositides, kinases, and GTPases^49^, including cdc42, RhoA^50^, RalA and RalB^51^. The exocyst is a Ral effector^51^, and active, GTP bound forms of both RalA and RalB can bind to Sec5 and Exo84^52^ and mediate the assembly of mature exocyst complexes from 4-protein precursor subcomplexes I and II^53^. We note that other proteins involved in exocytic vesicle fusion were identified as Merlin-proximal, including Rab11b and its effector WDR44, the v-SNAREs SNAP-29 and YKT6 and the syntaxin binding proteins STXBP1 and STXBP2, key regulators of the SNARE complex that is essential for exocytic vesicle fusion to the plasma membrane (Table II, Supplemental Spreadsheet 1). Indeed, SNAP-29 is one of the 7 proteins that, like RalB, is proximal to both Merlin isoforms only at confluence.

### Exocytosis in Cancer Biology

Fusion of exocytic vesicles with the plasma membrane is the mechanism by which integral membrane proteins and membrane lipids are incorporated into the plasma membrane and intra-vesicular contents are released into extracellular space^54^. The RalB-dependent changes in exocytosis kinetics seen upon Merlin loss may reflect distinct subtypes of exocytic events that can be visualized by the VAMP2-pHluorin reporter. Exocytic events with a longer duration are characteristic of “full fusion” exocytosis in which the vesicle membrane completely integrates with the plasma membrane, along with new membrane components and integral membrane proteins including cell junctional proteins and growth factor receptors^55^. Short duration exocytic events include “kiss-and-run” exocytosis, in which the vesicle opens and closes transiently, leading to the release of soluble intra-vesicular contents into the extracellular space, without incorporation of the vesicular membrane and associated integral membrane proteins into the plasma membrane^56^. The shift from shorter to longer duration events upon loss of Merlin is consistent with a shift from the “kiss and run” to the “full fusion” exocytic events that require the exocyst. Loss of exocyst function impairs “full fusion” and increases the proportion of “kiss and run” events^57^. Therefore, the changes in exocyst kinetics that we observed may reflect a shift to more “full fusion” events mediated by the exocyst in response to the increased RalB activity seen in Merlin KO cells at confluence or by the active RalB^G23V^ mutant in control cells.

Ral-regulation of exocytosis is crucial for cell growth, cell-cell communication, and establishment of cell polarity^58^. During development of highly polarized Schwann cells, Ral GTPases promote radial axonal sorting in peripheral nerves through the exocyst complex^59^. NF2 mutant schwannoma are epithelioid cells characterized by dysregulated polarity, caused by loss of Merlin function^60^. A key mechanism by which cell polarity is established and maintained is asymmetric membrane trafficking controlled by the exocyst^37^ and Ral activity is implicated in this process^61^. We hypothesize that upon contact-inhibition, cells are not actively growing and do not require plasma membrane expansion, full fusion exocytosis is down regulated by active-PIP_2_-bound Merlin dimers that bind to RalA/B and to prevent association with the exocyst and inhibit full fusion exocytosis. In NF2-schwannomatosis the absence of Merlin the failure to inhibit the RalA/B interaction with the exocyst causes increased “full fusion” exocytosis, leading to changes in the cell surface proteome that affects cell polarity in tumor cells.

### Therapeutic Implications

The identification of RalA and RalB as binding partners for active, PIP_2_-bound Merlin define a mechanism underlying previously described trafficking and cell polarity defects in *NF2* null cells. The coordinated nature of exocytosis and endocytosis and the correlation of increased exocytic duration with “full fusion” exocytosis^57^ raises the possibility that Merlin loss may cause differential expression of key proteins on the cell surface. The cell surface is the critical interface between tumor cells and the extracellular environment and is the point of interaction with pharmacological agents. Increased trafficking of differentially expressed cell surface proteins may also make these cells vulnerable to endocytic nanoparticle-based drug delivery strategies^62^. Constitutive RalB activity has been shown to be critical for survival in other human tumor cells^63^. This effect is mediated by Sec5-dependent activation of the IkB kinase family member TBK1^64^, raising the possibility that pharmacological inhibition of the Ral itself, or of its downstream effector pathways, may also induce death in schwannoma cells. The observation that loss of Merlin leads to increased RalA/B activity in confluent Schwann cells also identifies them as potential therapeutic target in NF2 schwannoma. As Ral-specific inhibitors for *in vivo* use become available, it will be of great interest to test their effects in this tumor type. That Ral small GTPases represent a new signal transduction pathway that may be exploited, either alone or in combination with agents targeting the HIPPO pathway, to treat NF2-related schwannomatosis.

## Materials and Methods

### Recombinant DNA

All recombinant DNA subcloning was performed using NEBuilder® HiFi DNA Assembly Cloning Kit (New England Biolabs, Ipswich, MA, Cat# E5520s). DNA fragments were PCR amplified using Q5 High-Fidelity 2X Master Mix (New England Biolabs, Ipswich, MA, Cat# M0492S) and purified Monarch® DNA Cleanup Columns (New England Biolabs, Ipswich, MA, Cat# T1034L). Oligonucleotide primers were purchased from Integrated DNA Technologies (Coralville, Iowa). The products of recombinant DNA manipulations were confirmed by sequencing (CCHMC Genomics Sequencing Facility, Cincinnati, OH). The pLNh-APEX vector was derived from plasmids lentiCRISPR, APEX2-GBP and pLenti-CMV-GFP using the NEBuilder HiFi DNA Assembly Cloning Kit (New England Biolabs, Cat# E5520s). The cDNAs for Merlin isoform1 and isoform 2 were cloned in pLNh-APEX in frame N-terminal to APEX2. pLenti-CMVie-XX-mCh was constructed by cloning the mCherry ORF from pcDNA3-FlipGFP(Casp3 cleavage seq) T2A mCherry into pLenti CMVie-IRES-BlastR. RalB was subcloned into the vector pLenti-CMVie-XX-mCh-BlastR in frame, C-terminal to mCherry to generate the lentiviral expression vector pLnCMV-Ch-RalB. The RalB mutants G23V, Q72L and D28N were constructed by PCR mediated site directed mutagenesis using primer containing the appropriate mutant. The vector pLentiCMV-VAMP2-pHluorin-Hygro was constructed by subcloning the VAMP2-pHluorin ORF in place of GFP in pLenti-CMV-GFP. The plasmids expressing GFP fusions for a-Catenin, d-catenin, PPP1R12A, RhoA, RALA and RALB cDNA ORF Clones cDNA ORF Clone, were purchased from Sino Biological US, Inc. (Wayne, PA, Cat#: HG-12388-ACG, HG-16488-ACG, HG-13990-ACG, HG12110-ANG, HG52476-ANG, HG15052-ANG). The plasmids pEGFP-C3-Exo70, pEGFP-C3-Sec8, pEGFP-C3-Exo84 and pEGFP-C3-Sec5 were a gift from Channing Der. (Addgene plasmids # 53761, # 53758, #53762, #53756. pClneoMyc human ERBIN was a gift from Yutaka Hata (Addgene plasmid # 40214. Full-length MLCK-GFP was a gift from Anne Bresnick (Addgene plasmid # 46316. pcDNA3-EGFP-Cdc42(wt) was a gift from Klaus Hahn (Addgene plasmid # 12599).

Cadherin-EGFP was a gift from Valeri Vasioukhin (Addgene plasmid # 18870. Integrin β1-GFP was a gift from Martin Humphries (Addgene plasmid # 69804). EGFP-Rab7A was a gift from Qing Zhong (Addgene plasmid # 28047). Human b-catenin GFP was a gift from Alpha Yap (Addgene plasmid # 71367). pEGFP-Akt1(WT) and pEGFP-Akt2 were a gift from Thomas Leonard & Ivan Yudushkin (Addgene plasmids # 86637, #86593. pLVX-EF1a-EGFP-RAB11B-IRES-Puromycin was a gift from David Andrews (Addgene plasmid # 134860). pLenti-PGK-RAP1A-WT-neo was a gift from Roland Friedel (Addgene plasmid # 156173). pCMV-Vamp2-pHluorin was a gift from Brad Zuchero (Addgene plasmid # 190151). The plasmids pEGFP-C3-Exo70, pEGFP-C3-Sec8, pEGFP-C3-Exo84 and pEGFP-C3-Sec5 were a gift from Channing Der. (Addgene plasmids # 53761, # 53758, #53762, #53756). pLenti CMVie-IRES-BlastR was a gift from Ghassan Mouneimne (Addgene plasmid # 119863). pcDNA3-FlipGFP(Casp3 cleavage seq) T2A mCherry was a gift from Xiaokun Shu (Addgene plasmid # 124428. The plasmid lentiCRISPR v2 was a gift from Feng Zhang (Addgene plasmid # 52961. APEX2-GBP was a gift from Rob Parton (Addgene plasmid # 67651). pLenti-CMV-GFP Hygro (656-4) was a gift from Eric Campeau & Paul Kaufman (Addgene plasmid # 17446. IKK1-EYFP was a gift from Johannes A. Schmid (Addgene plasmid # 111206). The plasmids mEmerald-Ezrin-N-14, mEmerald-Moesin-N-14, mEmerald-PTK2-C-10, mEmerald-ILK-N-17 were a gift from Michael Davidson (Addgene plasmid #54090, #54185, #54240, #54127). IKK1-EYFP was a gift from Johannes A. Schmid (Addgene plasmid # 111206). R777-E187 Hs.PPP1CA was a gift from Dominic Esposito (Addgene plasmid # 70471). Lenti-CAG-GFP-hKRAS4B-Zeocin was a gift from Dr. Liang Hu, (Cincinnati, Children’s Hospital). The lentiviral packaging plasmids psPAX2 and pMD2.G were a gift from Didier Trono (Addgene plasmids #12260, #12259)

### Cell Culture

The immortalized human Schwann cell line, iHSC1-λ, was gift from Dr. M. R. Wallace^24^. HEK-293T cells were purchased from ATCC (CRL-2316). Cells were grown in DMEM supplemented with 10% FBS and Pen/Strep at 37°C 5% CO_2_. 293T cells were transfected with 25000 MW, linear polyethylemimine (PEI, Polysciences, Warrington, PA. (cat# 23966-1). For lentiviral production, HEK-293T cells were co-transfected with the lentiviral expression plasmid and the packaging plasmids psPAX2 and pMD2.G using PEI (1.62 pmol:1.32 pmol:1.32 pmol ratio). After 24 hours, the media was changed and cells were grown in a minimal volume of fresh media for 48 hours. Conditioned media was collected, passed through a 0.22µ filter, aliquoted and stored at −80°C. iHSC-1λ cells were infected with a titration of lentiviral conditioned media in the presence of 8 µg/ml polybrene (MilliporeSigma, St. Louis, MO). After 48 hours, the cells were washed 3x with sterile PBS and grown in selection media, with either 2 µg/ml puromycin, 100 µg/ml hygromycin, or 5 µg/ml blasticidin.

### Immunofluorescence

Cells were plated in 8-well chamber slides (Cat# 80806-90, Ibidi, Fitchburg, WI). Cells were fixed in 4% paraformaldehyde, permeabilized, blocked and probed with primary antibodies, washed in PBS probed with Alexa^488^ or Alexa^555^ conjugated secondary antibodies (Cat# A32731, A32727, ThermoFisher Waltham, MA) and counterstained with DAPI and Alexa-647 conjugated-phalloidin (Cat# 8940S, Cell Signaling Technology, Danvers, MA). Images were acquired on a Nikon C2plus Confocal microscope using either a 20x, 0.75 NA, Nikon PlanApo λ or a 60x 1.4 NA, Apo λS Oil objective. Super-resolution microscopy was performed in the CCHMC Bio-Imaging and Analysis Facility (RRID:SCR_022628). Images were acquired on a Nikon Ti2, AX confocal equipped with an NSPARC ISM spatial array confocal detector in SR mode using a 100x, 1.45 NA, PLAN APO λD, Oil objective.

The antibodies used were: Merlin (Cat# 6995, Cell Signaling Technology, Davners, MA), RalA/B (Cat# sc-374582, Santa Cruz Biotechnology, Dallas, TX), Active Ral (Cat# 26913, NewEast Biosciences, Glenmoore, PA), RalA (Cat# 21034, NewEast Biosciences, Glenmoore, PA), RalB (Cat# 21033, NewEast Biosciences, Glenmoore, PA), YAP1 (Cat# PA1-46189, ThermoFisher, Waltham, MA), N-Cadherin (Cat# 14215, Cell Signaling Technology, Davners, MA), Exo70 (Cat# MABT186, Millipore Sigma, St Louis, MA), Exo70 (Cat# sc-365825, Santa Cruz Biotechnology, Dallas, TX), APEX (Cat# CAC09177, Biomatik, Kitchener, ON, Canada), Sec5 (Cat# sc-393230, Santa Cruz Biotechnology, Dallas, TX).

### Proximity Biotinylation

Three cell lines were generated by infecting iHSC-1λ cells with lentivirus expressing the fusion proteins Merlin isoform 1-APEX2, Merlin isoform 2-APEX2 or control APEX2. Hygromycin resistant cell populations were then selected and pooled. APEX activity was measured in lysates using ECL Chemiluminescent HRP substrate (Cat# WBKLS0050, Millipore Sigma, St Louis, MO). Proximity biotinylation was performed in triplicate in 15 cm dishes. For sub-confluent samples cells were plated at 5000 cells per cm^2^, confluent samples were plated at 15,000 cells per cm^2^. Cells were incubated for 48 hours then processed for biotinylation. Proximity biotinylation was performed as described in the literature^25^. Briefly, media was changed to fresh media supplemented with 500 µM biotin-tyramide (Cat#. SML2135-50MG, MilliporeSigma, St. Lousi, MO) and incubated for 30 min @ 37°C. To initiate biotinylation, fresh 100 mM H_2_O_2_ in PBS was added to 1 mM final concentration, cells were incubated at 37°C for 1 minute. The reaction was then quenched in final concentration of 5 mM Trolox (Cat# 238813-5g, MilliporeSigma, St Louis, MO), 10 mM sodium ascorbate and 10 mM sodium azide. Cells were then rinsed with ice cold PBS and lysed in RIPA buffer supplemented with a protease inhibitor cocktail (HALT, Pierce) and Benzonase (Sigma-Aldrich). Biotinylated proteins were isolated from cleared lysates by incubating with Streptavidin-Mag agarose beads (Cat# GE28-9857-99, MilliporeSigma, St Louis, MO) overnight at 4°C with agitation. Beads were captured in a magnetic stand, washed 3x with RIPA, 3x with 2M urea in TBS pH 7.4, 3x with TBS pH 7.4 and 5% aliquots were resuspended in a fluorescent protein staining loading buffer (Instant Bands, Cat# PFS001P, Thomas Scientific, Swedesboro, NJ) run on 4-20% gradient SDS–polyacrylamide gel electrophoresis (PAGE) (Bio-Rad). Fluorescent total proteins were visualized in an Azure C-600 imaging system (Azure Biosystems, Dublin, CA). Gels were then blotted and probed with streptavidin-IRDye680 and antibodies to Merlin. The remaining beads were washed 10x with 50 mM ammonium bicarbonate then sent to the Taplin Biological Mass Spectrometry Facility at Harvard Medical School and processed for matrix-assisted laser desorption/ionization mass spectroscopy. Mass spec data consisting of peptide number mapping to specific proteins were normalized to the total number peptides per sample, expressed per 10,000 peptides. Data was displayed as a proximity network using Cytoscape^65^. Gene ontogeny and functional enrichment was performed within Cytoscape using the String database ^312^. Hierarchical cluster analysis was performed using the web application Morpheus (Broad Institute, MIT, https://software.broadinstitute.org/morpheus).

### Statistics

Collated mass spec data was analyzed for statistically significant increases over the APEX only control by multiple unpaired T-test and a 5% false discovery rate using the two-stage step-up method^66^ with Graphpad Prism 10. For binding assays, unpaired T-test were used to test significance.

### Merlin Binding Assays

Direct Merlin binding assays were performed, with modifications, as previously described^11,14^. Assays were performed in triplicate, by incubating a probe consisting of purified Merlin-NanoLuc fusion protein with magnetic beads coated with GFP-fused bait proteins, isolated from transiently transfected 293T cells, in the presence or absence of 200 µM PI(4,5)P_2_ diC8 (Cat# P-4058-2, Echelon Biosciences, Salt Lake City, UT).

### Merlin NanoLuc Probe Purification

Twelve 15 cm dishes with 1.3 × 10^7^ 293T cells per plate were transfected with plasmid expressing Merlin fused N-terminal to NanoLuc with a dual Streptag affinity tag at its C-terminus (pMerNLst2) using 1 mg/ml PEI, at a 5:1 µl PEI to µg DNA ratio (250 µl PEI to 50 µg plasmid, per plate). Cells were incubated for 48 hours. Cell lysates were harvested by washing with ice cold PBS, then lysing in a high salt lysis buffer (20 mM TrisCl pH 8.0, 500 mM NaCl, 10 mM MgCl_2_, 0.5% NP-40 supplemented with 1x HALT protease and phosphatase inhibitor, Cat# 78430, ThermoFisher, Waltham, MA) and 1:5000 Benzonase (Cat# E1014-5KU, Millipore Sigma, St Louis, MA). Lysates were cleared by centrifugation at 21,000 xg for 10 min. at 4°C. Merlin-NanoLucST2 protein was affinity purified by gravity flow over a 0.2 ml bed volume Strep-TactinXT 4Flow high-capacity resin column (Cat# 2-5030-002, ThermoFisher, Waltham, MA). Columns were washed with 15 column volumes (CV) High Salt Wash Buffer (500 mM NaCl, 20 mM Tris-Cl pH 8.0, 0.1% Triton X-100, 0.1% Tween-20 + HALT), then 15 CV Normal Salt Wash Buffer (150 mM NaCl, 20 mM Tris-Cl pH 8.0, 0.1% Triton X-100, 0.1% Tween-20 + HALT) followed by 15 CV of TBS (150 mM NaCl, 20 mM Tris-Cl pH 8.0). Merlin-NanoLucST2 was eluted stepwise with 0.3 ml of Elution Buffer (100 mM Tris-Cl pH 8.0, 150 mM NaCl, 50 mM Biotin + HALT) per step for a total of 8 fractions. Purification was monitored by NanoLuc luciferase assays on all fractions (Cat# N1110, Promega, Madison WI), measured using white round bottom 96-well plates in a FlexStation 3 plate reader (Molecular Devices, San Jose, CA). Positive fractions were pooled, passed through a 0.22 µ filter then loaded onto a NCG Quest plus FPLC for gel filtration over a 10×300 Enrich SEC 650 size exclusion column (Bio-Rad, Hercules, CA) equilbtated in TBS (20 mM Tris-Cl pH 7.4, 150 mM NaCl).

Collected fractions were assayed for NanoLuc activity. High activity fractions corresponding to the closed conformation^11^ eluting at 13-14 ml were pooled. Concentration was determined by A^280^ in a Nanodrop 2000c (ThermoFisher, Waltham, MA). Purity was determined by SDS-PAGE using Instant Bands loading dye imaged on the Azure C-600 imaging system (Azure Biosystems, Dublin, CA). Preps were adjusted 0.5 mg/ml BSA and 0.1% Tween-20 and stored at 4°C for no more than two weeks.

### GFP-Bait Protein Purification

GFP-bait expressing plasmids were transfected into 5 × 10^6^ HEK 293T cells in a 10 cm dish and incubated for 48 hours. Cells were rinsed with cold PBS, harvested in Lysis Buffer (10 mM Tris-Cl pH 7.5, 150 mM NaCl, 2 mM, MgCl_2,_ 0.5 mM EDTA, 0.5% NP-40 + 1x HALT and 1:5000 Bensonase), cleared by centrifugation at 21,000 xg for 10 min. at 4°C. Lystates were incubated with 30 µl washed GFP-Trap_MA magnetic agarose beads (Cat# GTMA020, Bulldog Bio Inc. Portsmouth, NH) overnight at 4°C with agitation. The beads were captured, washed three times with High Salt Wash Buffer (20 mM Tris-Cl pH 7.4, 0.5 M NaCl, 0.1% Triton X-100, 0.1% Tween 20), followed by three washes with Low Salt Wash Buffer (20 mM Tris-Cl pH 7.4, 50 mM NaCl, 10% Glycerol, 0.2% NP-40) then three washes with Normal Salt Wash Buffer (20 mM Tris-Cl pH 7.4,150 mM NaCl, 0.1% Triton X-100, 0.1% Tween 20) and finally resuspended in Blocking Buffer (20 mM Tris-Cl pH, 7.4, 150 mM NaCl, 2.5 mg/ml BSA, 0.05% Tween-20). Final bead fluorescence was measured on a FlexStation 3 plate reader (Molecular Devices, San Jose, CA).

### Direct Binding Assays

Each preparation GFP-bait-bound was divided into six wells of a white round bottom 96-well plates. The beads were captured on a 96-well magnetic stand (EpiMag HT (96-Well) Magnetic Separator, Cat# Q10002-1, EpiGenTek, Farmingdale, NY), resuspended in a 30-µl Blocking Buffer and incubated for 1 hour at room temperature with agitation. The beads were recovered magnetically then resuspended in 100 nM Merlin-NLuc protein in Blocking Buffer with or without 200 µM PI(4,5)P_2_ diC8 and incubated at room temperature for 1 hour with agitation. Beads were then captured magnetically, washed four times with TBST, resuspended in 25 µl TBST and NanoLuc luciferase activity was measured using NanoGlo Luciferase Substrate Buffer (Promega, Madison, WI). NanoLuc activity, GFP fluorescence and BRET spectra were measured on a FlexStation 3 (Molecular Devices, San Jose, CA). For display purposes the plates were also imaged on that Azure C-600 imaging system (Azure Biosystems, Dublin, CA). Data was expressed a % NanoLuc activity bound to the beads relative to the input Merlin-NanoLuc probe, normalized to the GFP-only negative control.

### Ral Activity and Exocytosis Assays

Ral activity was assayed in sub-confluent and confluent Control and Merlin knockout iHSC-1λ cells using the RalB activity assay (Cat# 81501, NewEast Biosciences, Glenmoore, PA) by immunoprecipitation with antibodies to active Ral and then probes with antibodies specific for RalA and RalB, (Cat # 26913, 21034, 21033, NewEast Biosciences, Glenmoore, PA). For exocytosis assays, Control and Merlin knockout iHSC-1λ cells were infected with lentivirus expressing VAMP2-pHluorin and selected with 100 µg/ml hygromycin. Subsequently, both resulting cell lines were infected with lentivirus expressing mCherry-RalB^wt^, mCherry-RalB^G23V^, mCherry-RalB^D28N^ and control mCherry and expressing clones were selected with 5 µg/ml blasticidin. Exocytosis was measured in live VAMP2-pHluorin expressing cells in the CCHMC Bio-Imaging and Analysis Facility (RRID:SCR_022628). Imaging was performed on a TIRF equipped Nikon TiE microscope equipped with a Tokai Hit stage top incubation system and an Andor DU-897 C-6347 EMCCD camera using a Nikon Apo TIRF 100x Oil DIC, 1.49 NA. Time lapse images of live VAMP2-pHluorin expressing cells were acquired using TIRF illumination for 2 min. at 100 msec intervals. Exocytosis was analyzed using the ExoJ plugin^67^ on FIJI/Image J software run on the CCHMC High-Performance Computing cluster(RRID:SCR_022622).

## Supporting information

Collated Merlin Proximal Mass Spec Data

GO Analysis

## Acknowlegements

This work was supported by the Department of Defense, Congressionally Directed Medical Research Program Neurofibromatosis Research Program Awards NF120118, NF160078 and NF190083 and the University of Cincinnati Brain Tumor Center Jejurikar Fellowship Program to RFH. We thank Dr. Shyra Tedesco and Dr. Brad Ozanne for critically reading the manuscript. We would like to acknowledge the support of the CCHMC Bio-Imaging and Analysis Facility (RRID:SCR_022628) and the Information Services for Research (IS4R) (RRID:SCR_022622) shared facilities at Cincinnati Children’s Hospital Medical Center.

## Notes

### Competing Interest Statement

The authors have declared no competing interest.

