## Supplemental Figures for "Active Merlin Binds RalB to Regulate Exocytosis"

Supplemental Figure 1. Generation of Merlin knockout iHSC-1λ Schwann cells.

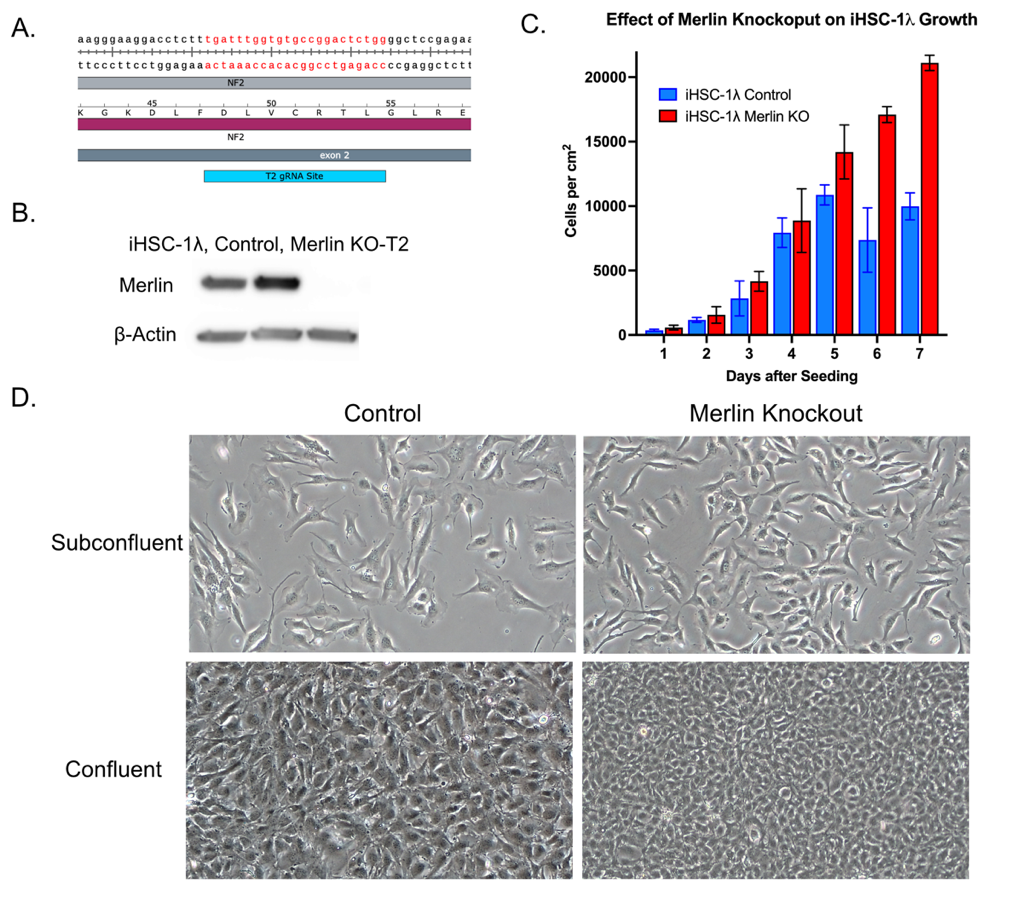

Supplemental Figure 1. Generation of Merlin knockout iHSC-1λ Schwann cells.

A. A graphical depiction of the gRNA target site in exon 2 of the human NF2 gene on Chromosome 22.
B. Immunoblot analysis of lysates from parental iHSC-1λ Schwann cells, control cells with a randomly scrambled gRNA and Merlin knockout cells probed with antibodies to Merlin (top) and β-actin (bottom) as a loading control.
C. Seven-day growth curves of Control (blue) and Merlin knockout (red) cells, showing a roughly two-fold increased cell density at confluence for Merlin knockout relative to Control cells.
D. Phase contrast images of Control (left panels) and Merlin knockout (right panels) cells in subconfluent (top) and confluent (bottom) cell densities. Note the increased saturation density evident in Merlin knockout cells relative to controls at confluence.

Supplemental Figure 2. Contact Inhibition.

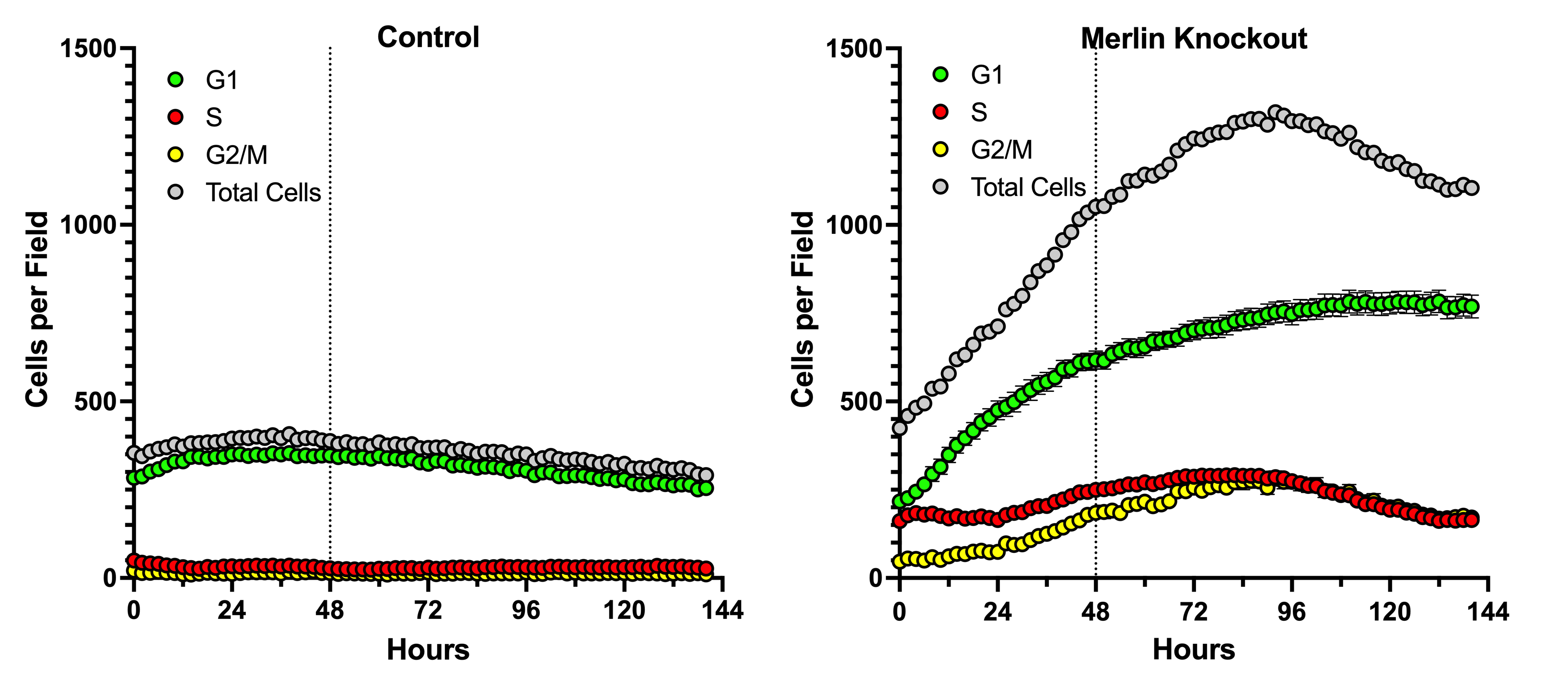

Supplemental Figure 2. Contact Inhibition.

Cell cycle dynamics of contact inhibition of growth was measured in cells expressing the PIP-FUCCI cell cycle reporter for control (left panel) and Merlin knockout (right panel) cells. Cells were plated at high cell density (~15,000 cells per cm^2^) in triplicate and G1 phase (green only), S-phase (red only) and G2/M phase (yellow) were counted every two hours for 6 days using an IncuCyte live cell imaging system. Total cells are the sum of G1-, S- and G2/M-phase cells and are depicted in grey. The dotted line at 48 hours indicated the harvest point for the proximity biotinylation experiments shown in Figure 2.

Supplemental Figure 3 Subcellular Distribution of Merlin Proximal Cells.

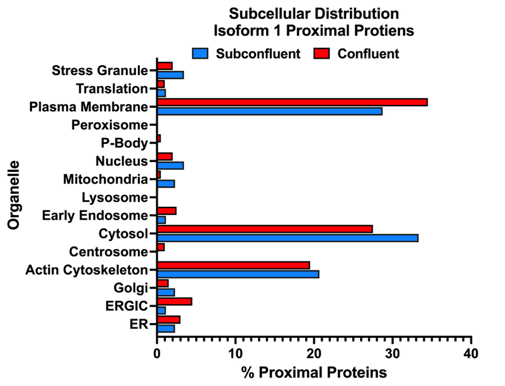

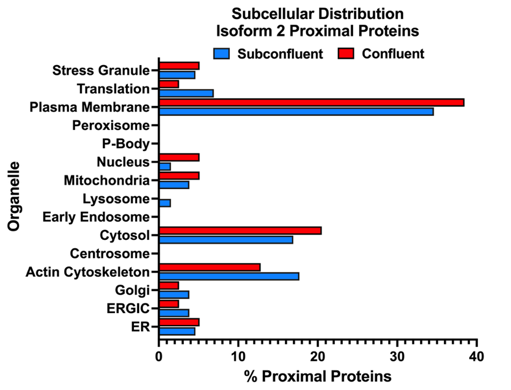

Supplemental Figure 3 Subcellular Distribution of Merlin Proximal Cells.

Subcellular distribution of Merlin proximal proteins displayed as a percentage of the total proximal proteins for both Merlin isoform 1 (left) and isoform 2 (right) in subconfluent (blue) and confluent (red) cells.
